# Extracellular matrix phenotyping by imaging mass cytometry defines distinct cellular matrix environments associated with allergic airway inflammation

**DOI:** 10.1101/2024.11.15.623782

**Authors:** J E Parkinson, M Ghafoor, R J Dodd, H E Tompkins, M Fergie, M Rattray, J E Allen, T E Sutherland

## Abstract

The extracellular matrix (ECM) forms the scaffold in which cells reside and interact. The composition of this scaffold guides the development of local immune responses and tissue function. With the advent of multiplexed spatial imaging methodologies, investigating the intricacies of cellular spatial organisation are more accessible than ever. However, the relationship between cellular organisation and ECM composition has been broadly overlooked. Using imaging mass cytometry, we investigated the association between cellular niches and their surrounding matrix environment during allergic airway inflammation in two commonly used mouse strains. By first classifying cells according to their canonical intracellular markers and then by developing a novel analysis pipeline to independently characterise a cells ECM environment, we integrated analysis of both intracellular and extracellular data. Applying this methodology to three distinct tissue regions we reveal disparate and restricted responses. Recruited neutrophils were dispersed within the alveolar parenchyma, alongside a loss of alveolar type I cells and an expansion of alveolar type II cells. This activated parenchyma was associated with increased proximity to hyaluronan and chondroitin sulphate. In contrast, infiltrating CD11b^+^ and MHCII^+^ cells accumulated in the adventitial cuff and aligned with an expansion of the subepithelial layer. This expanded subepithelial region was enriched for closely interacting stromal and CD11b^+^ immune cells which overlaid regions enriched for type-I and type-III collagen. The cell-cell and cell-matrix interactions identified here will provide a greater understanding of the mechanisms and regulation of allergic disease progression across different inbred mouse strains and provide specific pathways to target aspects of remodelling during allergic pathology.

**Graphical Abstract:** 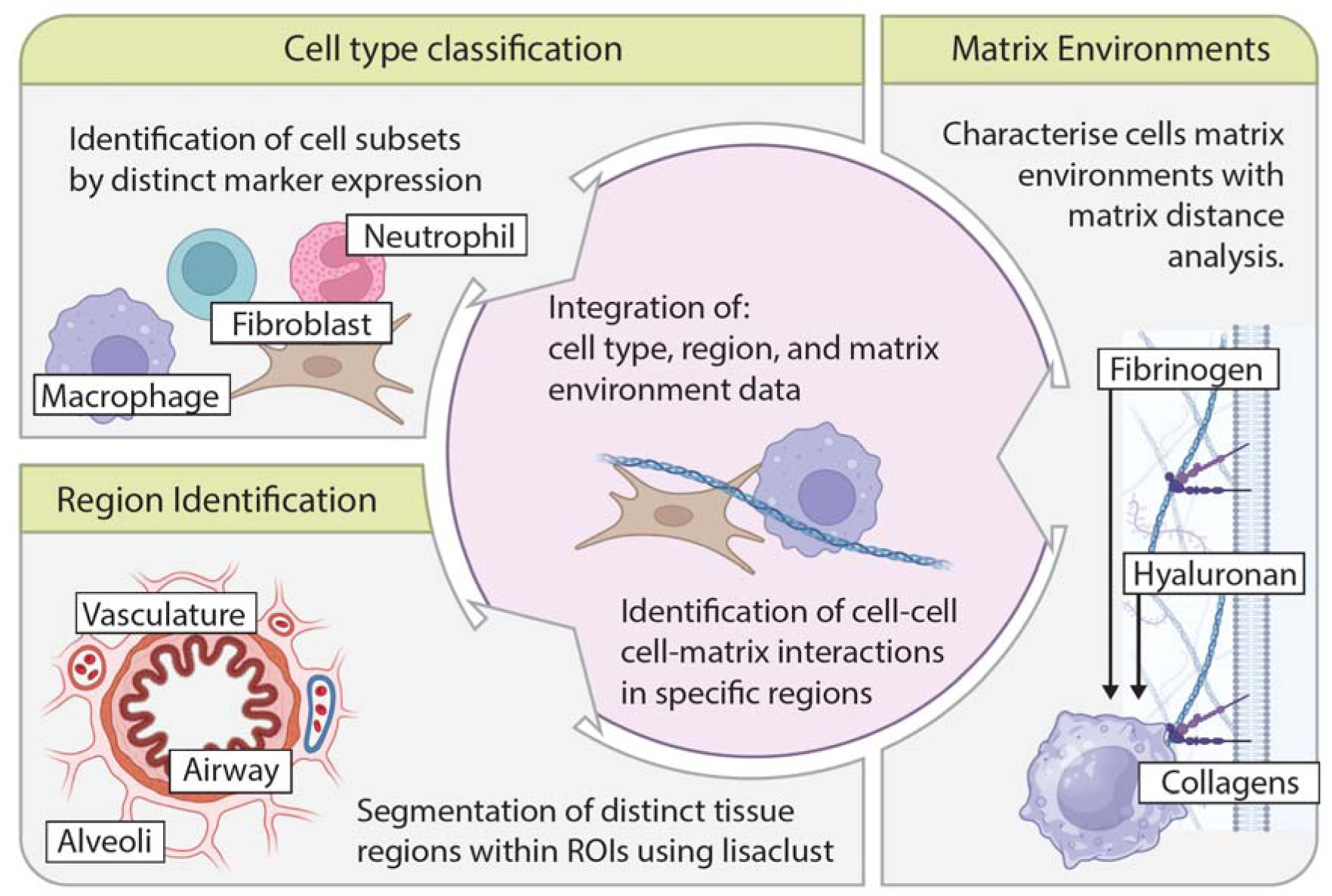

## Introduction

The extracellular matrix (ECM) is a highly heterogenous structure which comprises a vast array of molecules including proteins, glycosaminoglycans, and proteoglycans. Some of the most widely studied components are collagens, but the ECM also contains, among others: laminins, fibronectin, hyaluronan, and a diverse array of associated proteins^1^. These components are known to be organised into complex and distinct networks which confer important properties to the tissue, such as stiffness and elasticity, which then influence cellular responses^2–6^. In addition to being the framework within which cellular interactions occur^7^, the ECM is also an important transducer of cellular signals^8^. Many of the immune system’s canonical signalling pathways are shaped by both direct and indirect interactions with the ECM^9^. Therefore, it is vital to acknowledge the mutual roles of cellular ECM environments when studying a tissue’s response to challenges such as injury or disease.

Alterations in the composition and organisation of the ECM are present in almost all diseases, though to different extents^10–16^. In respiratory diseases such as asthma, airway remodelling encapsulates a variety of processes including epithelial activation, smooth muscle hyperplasia, immune infiltration, and airway subepithelial ECM reorganisation^17,18^. Together, these features are a major driver of airway hyperresponsiveness and therefore, breathing difficulties in people with asthma^19^. To examine the processes that drive airway remodelling, we have utilised a DRA model, involving chronic exposure to extracts of three clinically relevant allergens including house dust mite, ragweed, and aspergillus^20^. During DRA induced allergic airway inflammation (AAI), deposition of type I and III collagens occurs alongside the pulmonary recruitment of effector inflammatory cells such as eosinophils and neutrophils^21^. The combination of ECM reorganisation and immune cell infiltration and activation makes AAI an ideal model for examining how the composition and organisation of the ECM correlates with inflammatory cell recruitment and disease progression.

With the advent of multiplexed imaging systems such as imaging mass cytometry (IMC)^22^, cyclic immunofluorescence (IF)^23–25^, and spatial transcriptomics^26^, tools now exist to quantify the spatial environment of different organs and disease systems in greater detail than ever before. Multiparameter flow cytometry, and single cell ribonucleic acid sequencing techniques have facilitated in-depth characterisation of the inflammatory cellular mileu^27^. However, these techniques rely on tissue dissociation and therefore fail to capture spatial information within the tissue. Whilst interrogating multiple markers spatially is traditionally done using IF staining, this is often limited to four or five targets due to spectral overlap between fluorophores^28^. Highly multiplexed tissue imaging studies have now been widely implemented either alone or integrated with transcriptomics and high-parameter flow cytometry. This has been applied particularly to the understanding of cell-cell interactions within the tumour microenvironment^22,29–32^. Recently this has been expanded to include other aspects of the immune response such as quantifying the cytokine milieu and how this changes across different tumour regions^33^. Despite these advances, relatively few studies have included an in-depth classification of the ECM environment, and even fewer studies have integrated ECM markers with cellular analysis.

Using the DRA mouse model of AAI^20^ we aimed to characterise cell-cell and cell-ECM interactions that may be present during allergic pathology. BALB/c and C57BL/6 mice were used to further interrogate links between inflammation and pathology, as these mouse strains have been shown to differentially regulate specific ECM components in this model^21^. IMC was employed with a new approach to integrate ECM analysis with existing multiplex image analysis pipelines. Data demonstrated that during AAI we can identify distinct matrix environments that associate with observable tissue structures, such as airways. The immune cell recruitment observed in DRA was localised within distinct lung adventitial or alveolar niches and these correlated with specific matrix environments in BALB/c and C57BL/6 mice. Furthermore, expansion of cells within the remodelled subepithelial space during AAI correlated with enhanced interactions between CD11b+ cells and stromal populations. Together, these data suggest that immune-stromal cell networks may form an integral component that drives and maintains a localised pathogenic extracellular matrix.

## Results

### Identification of cellular and extracellular molecules using IMC

There is a long publishing history examining pulmonary cellular changes during AAI^34–41^ however, very little work has examined the cellular composition in relation to the ECM. We sought to characterise the spatial lung organisation in BALB/c and C57BL/6 mice after chronic administration of PBS (control) or an allergen cocktail containing house dust mite, ragweed, and aspergillus (DRA)^20,21^ (**Fig 1 a**). The lung environment was characterised using an IMC panel of 32 antibodies, including 19 populations markers, 3 activation markers, and 10 ECM markers (**Fig 1 b**). Markers were chosen to identify major epithelial, mesenchymal, and immune populations in the naïve and allergic lung, in addition to providing an overview of the surrounding ECM (**Fig 1 b**). To account for heterogeneity across the lung, three regions of interest (ROIs) per mouse were imaged across different lung lobes (**Fig 1 c**). ROIs encompassed small-medium airways, associated blood vessels, and the immediate alveolar parenchyma. Consistent with previous findings^21^, staining demonstrated immune cell recruitment following DRA administration in both mouse strains as well as potential alterations in airway epithelial cell (AEC) morphology (**Fig 1 d**).

**Figure 1:**
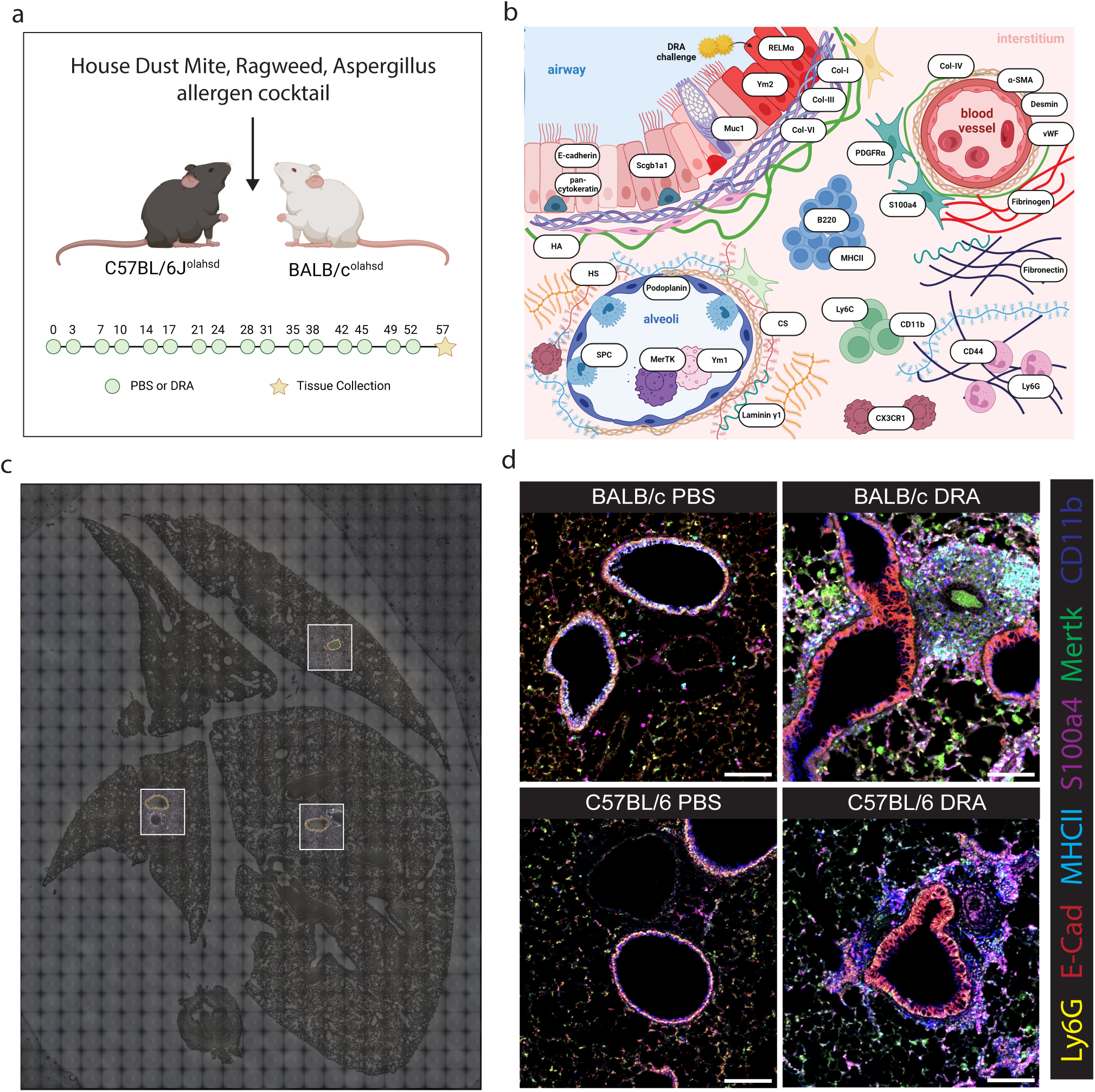
Identification of cellular and ECM molecules using IMC To investigate the association between the cellular and extracellular responses we utilised Hyperion multiplexed imaging analysis in a model of allergic airway inflammation. **a**) Schematic showing the experimental setup and dosing protocol for allergic airway inflammation. BALB/c and C57BL/6 mice were dosed twice weekly with a house <u>d</u>ust mite, ragweed, and <u>a</u>spergillus (DRA) combination allergen cocktail for eight weeks. **b**) A Hyperion® panel was designed to cover a wide range of cell populations as well as extracellular matrix components found across sub-structures within the lung, such as airways, vessels and alveoli. **c**) Representative images of IMC-stained tissue from a BALB/c PBS treated animal with the three ROIs highlighted in white. Areas were chosen to include at least one small-medium airway and associated structures. **d**) Images of antibody staining data showing a subset of cellular markers within a representative ROI from C57BL/6 and BALB/c mice administered PBS or DRA. Scale bar is equivalent to 160 μm. Panel **b** was generated with biorender.com.

### Distinct epithelial, stromal, and immune niches exist in the naïve and allergic lung

Raw data from each ROI was analysed using the steinbock pipeline^42^. Cell masks were created using deepcell^43^ and single cell expression data generated from the average intensity per channel within each object in the cell mask, calculated using the ski mage package^44^. Relative cell density, quantified by the percentage of the ROI covered by the cell mask, was increased during AAI (**Fig 2 a**). Cell density corresponded with increased total cell number, both of which were significantly greater in allergic BALB/c compared to allergic C57BL/6 mice (**Fig 2 b**).

**Figure 2:**
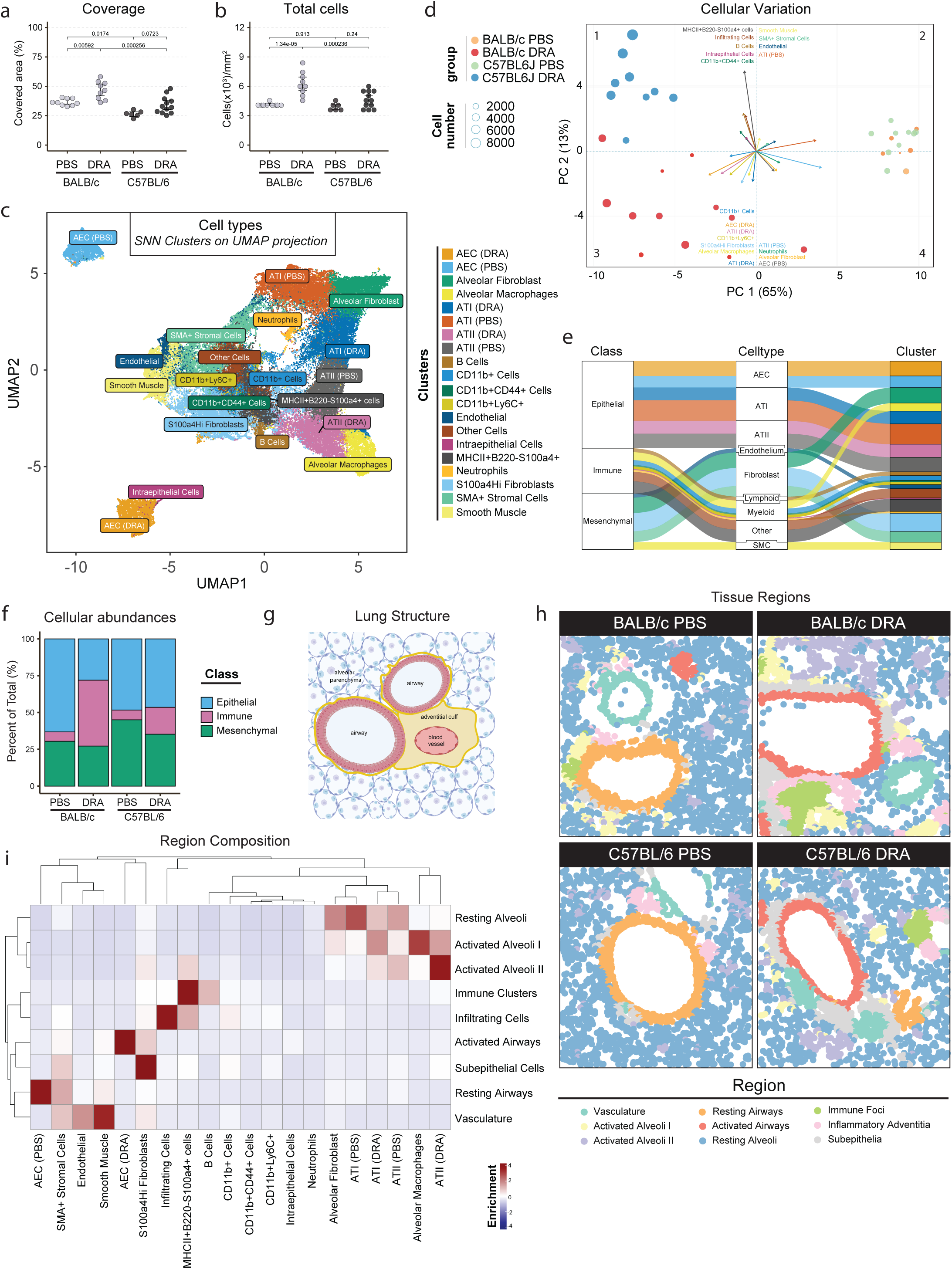
Lung regions are associated with distinct ECM environments that change during AAI. To define the cell populations within our ROIs cell masks were generated using deepcell and a single cell expression profile for each masked cell was quantified using scikit. **a**) Percentage of each ROI covered by the mask and **b**) total number of cells within each ROI. **c**) To group similar cells dimensionality reduction was performed using UMAP and distinct populations were grouped using SNN clustering followed by manual separation based on specific cellular expression data. **d**) Total variation within the dataset was visualised using centred log-ratios (CLR) on the cluster proportions across samples, shown on PCA reduction. Arrows represent loadings and are colour coded to cell type. **e**) Schematic showing the categorisation and relationship between different cell populations. **f**) To broadly characterise the response across mice, the abundance of cells in defined categories from **e** were quantified. **g**) Diagram showing the distinct structure of the lung including the main macro tissue structures. **h**) lisaClust was used to define spatially adjacent groups of cell clusters from **c** which was projected onto deepcell generated cell masks. **i**) The contribution of specific cell clusters to different lisaClust regions. Scale represents proportions of cell types making up each tissue region which have been scaled and centred relative to the row. For **a,b,f** data points represent individual ROIs with error bars showing the 95% confidence intervals around the mean (black filled circle). n = 2 - 4 animals per group corresponding to 6 - 12 ROIs per group. Statistical significance was calculated using a one-way ANOVA with Tukey post hoc comparison test. Panel **g** was generated with biorender.com.

To identify distinct cell populations in the lung, dimensionality reduction of the cellular staining was performed. Shared nearest neighbour (SNN) clustering^45^, followed by manual separation based on canonical marker expression, identified twenty clusters that were projected onto two dimensions using uniform manifold approximation and projection (UMAP) (**Fig 2 c**). Cells within the same cluster occupied similar UMAP space suggesting robust identification of cell populations. Identified clusters were annotated by their distinct marker expression patterns (**sup Fig 2 a**) and spatial localisation. Of note, SNN clustering split some populations, e.g. airway epithelial cells, into two clusters based on PBS or DRA treatment (**Fig 2 c** and **sup Fig 2 a**). Additional multidimensional scaling (MDS) showed similar cell types were found close together, such as different immune cells localised together in the middle of the plot (**sup Fig 2 b**). Principal component analysis and MDS clustered samples from different groups together and identified key cellular drivers that characterised allergic responses, including CD11b^+^ immune cells and S100a4^+^ fibroblast populations, among others (**Fig 2 d and sup Fig 2 c**). Interestingly, cellular drivers appeared to be distinct between BALB/c and C57BL/6 animals, but only under allergic conditions (**Fig 2 d**), again suggesting divergence in the response to DRA challenge between strains^21^. To simplify further analysis, identified cell clusters were classified based on their respective lineage and function into cell types and classes (**Fig 2 e**). Quantification of the proportion of different cell classes revealed a shift from predominantly pulmonary epithelial cells in PBS treated animals to an increased proportion of immune cells after DRA administration (**Fig 2 f**). Immune cell expansion was especially evident in BALB/c animals. Of note mesenchymal cells were more abundant in C57BL/6 PBS animals than their BALB/c PBS counterparts suggesting possible cellular differences between these strains in the steady state (**Fig 2 f)**.

Specific lung cell populations are associated with distinct tissue structures (**Fig 2g**) that carry out essential functions. Thus, we assessed whether organised tissue regions could be identified based on their constituent cell populations. The lisaClust^46^ algorithm was employed to define closely interacting groups of cells based on phenotypic classification alongside spatial localisation. Nine distinct spatial regions were identified across PBS and DRA ROIs (**Fig 2 h** and **sup Fig 2 d**). These regions were annotated according to their constituent cell types and localisation (**Fig 2 i**). Identified tissue regions included vasculature, alveolar epithelium, airway epithelium, subepithelial regions, inflammatory adventitia and immune foci. Several of these regions, including airway and alveolar epithelium, separated between PBS and DRA treated animals independently of strain, indicating differences in cell activation (**Fig 2 i**). Additionally, several regions associated with inflammatory cell recruitment were expanded in DRA challenged animals.

### Lung regions are associated with distinct ECM environments that change during AAI

Having identified distinct cell types and tissue niches within our dataset, we next asked how these niches related to ECM components included in our staining panel. These canonically extra-cellular molecules are not well represented by cell mask-based analysis (**Fig 3 a**). Hence, we developed a novel deep-learning thresholding algorithm (Deepthresh) combined with euclidian distance calculation to determine a cells distance to specific matrix molecules (**sup Fig 3 a, b,** and **methods**). To then classify similar cellular ECM environments, cells were grouped into clusters using SNN clustering of these matrix distances. This generated 12 lung matrix environments which were projected on to a UMAP (**Fig 3 b**) and showed distinct spatial patterns throughout the tissue and correlating with known structures such as airways, vessels, or alveolar epithelium (**Fig 3 c**). Matrix environments were annotated according to their nearest ECM components (**sup Fig 3 c**) and spatial localisation (**Fig 3 c**). Annotated environments were then correlated to previously generated lisaClust tissue regions (**Fig 3 d and Fig 2 h**). Two major compartments were evident within the lung structure. Firstly, the alveolar parenchyma, which was associated with heparan sulphate, laminin gamma-1, type-IV collagen, and fibrinogen. Secondly, the adventitial cuff surrounding the airways and blood vessels which was associated with type-I, III and VI collagens as well as hyaluronan (**Fig 3 d**). Together, our methodology successfully characterised matrix environments across the lung tissue.

**Figure 3:**
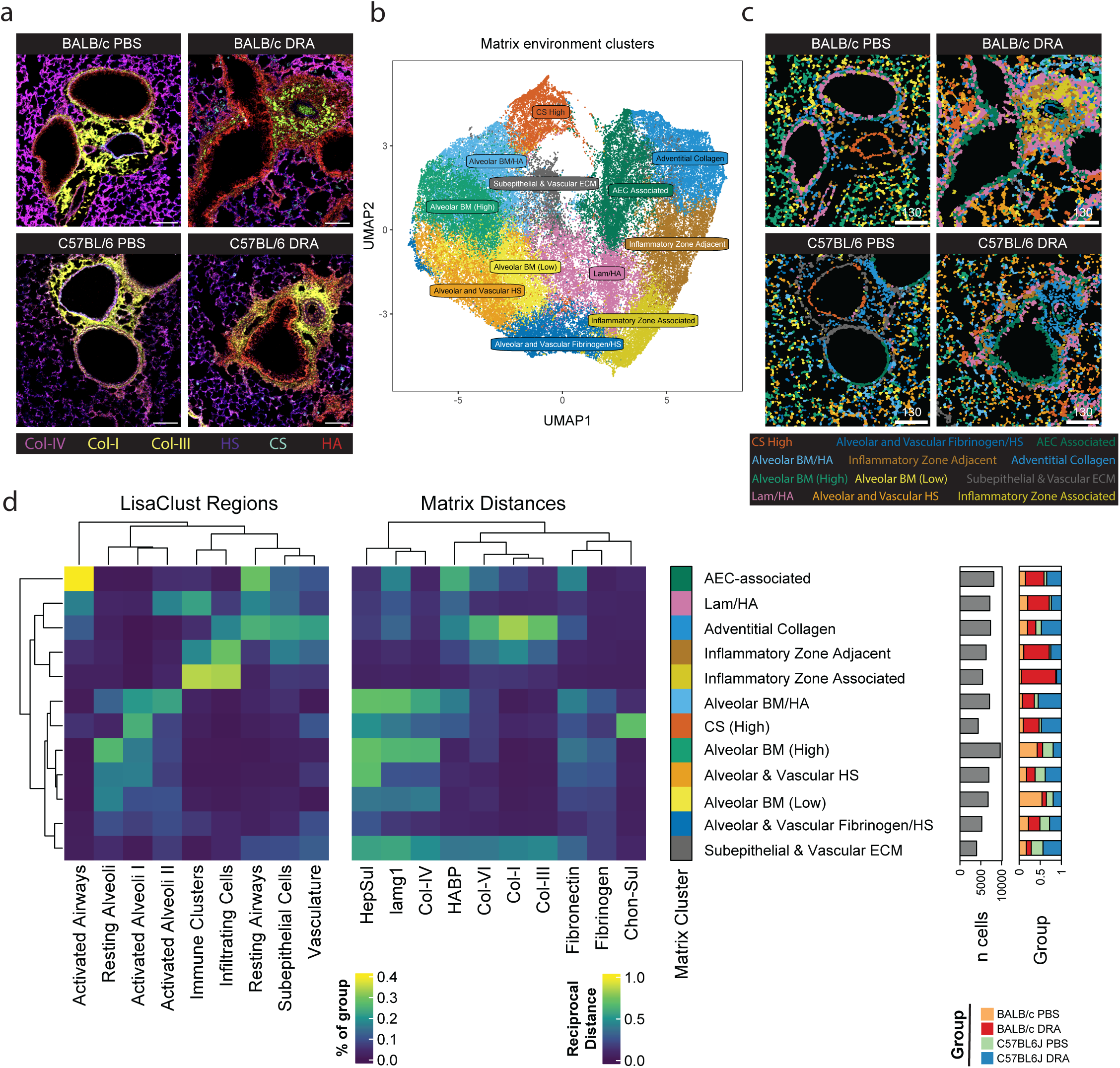
Lung regions are associated with distinct ECM environments which change during AAI. In order to determine if specific matrix components were associated with specific populations or treatments we aimed to characterise matrix environments of cell clusters. **a**) Example staining of several ECM molecules on representative ROIs. Matrix distances were calculated as described in the methods and then **b**) dimensionality reduction was performed using UMAP and distinct populations were grouped using SNN clustering to define distinct matrix environments. **c**) Projection of defined matrix environments onto deepcell generated cell masks for representative images. Scale bar equivalent to 130μm. **d**) To visualise the enrichment of specific matrix environments contingency tables were generated and plotted via heatmap. Left panel shows the percentage of cells in that region associated with the specific matrix environment. Right panel shows the average reciprocal distances to matrix components for specific matrix environments. Grey bars show the total number of cells assigned to different matrix environments and stacked bars show the relative contribution to different genotypes and treatments.

### Infiltrating immune cells localise in patches around the AAI adventitial cuff

Having established that tissue niches and matrix environments could be identified in our data, we next asked whether these changed in response to AAI in both C57BL/6 and BALB/c mice. AAI is traditionally associated with type-2 and type-17 immune responses and granulocyte recruitment^47,48^. Immune cells were detected in most tissue niches within ROIs, but we found that they were most enriched within the adventitial tissue, and this was particularly evident in BALB/c DRA-treated animals (**Fig 4 a**). This enrichment of immune cells correlated with the lisaClust defined inflammatory adventitia and immune foci regions (**Fig 4 b**). The Inflammatory adventitia consisted mainly of CD11b^+^ cells, whilst the immune foci were enriched for B220^+^ B cells (**Fig 2 i** and **Fig 4 c**). These regions were rarely observed in PBS-treated mice and only significantly expanded in BALB/c animals following DRA challenge (**Fig 4 d** and **Fig 4 e**). Aligning with this, both CD11b^+^ (**Fig 4 f**) and B220^+^ B cells (**Fig 4 g**) were further increased specifically in BALB/c DRA animals. This was also true for CD11b^+^CD44^+^ cells, infiltrating cells, and MHCII^+^B220^-^S100a4^+^ (**sup Fig 4 a-c**). However, other cell populations within the same tissue region, such as CD11b^+^Ly6C^+^ cells, expanded equivalently following DRA treatment in both strains, suggesting a separation in the response between certain cell types (**sup Fig 4 d**).

**Figure 4:**
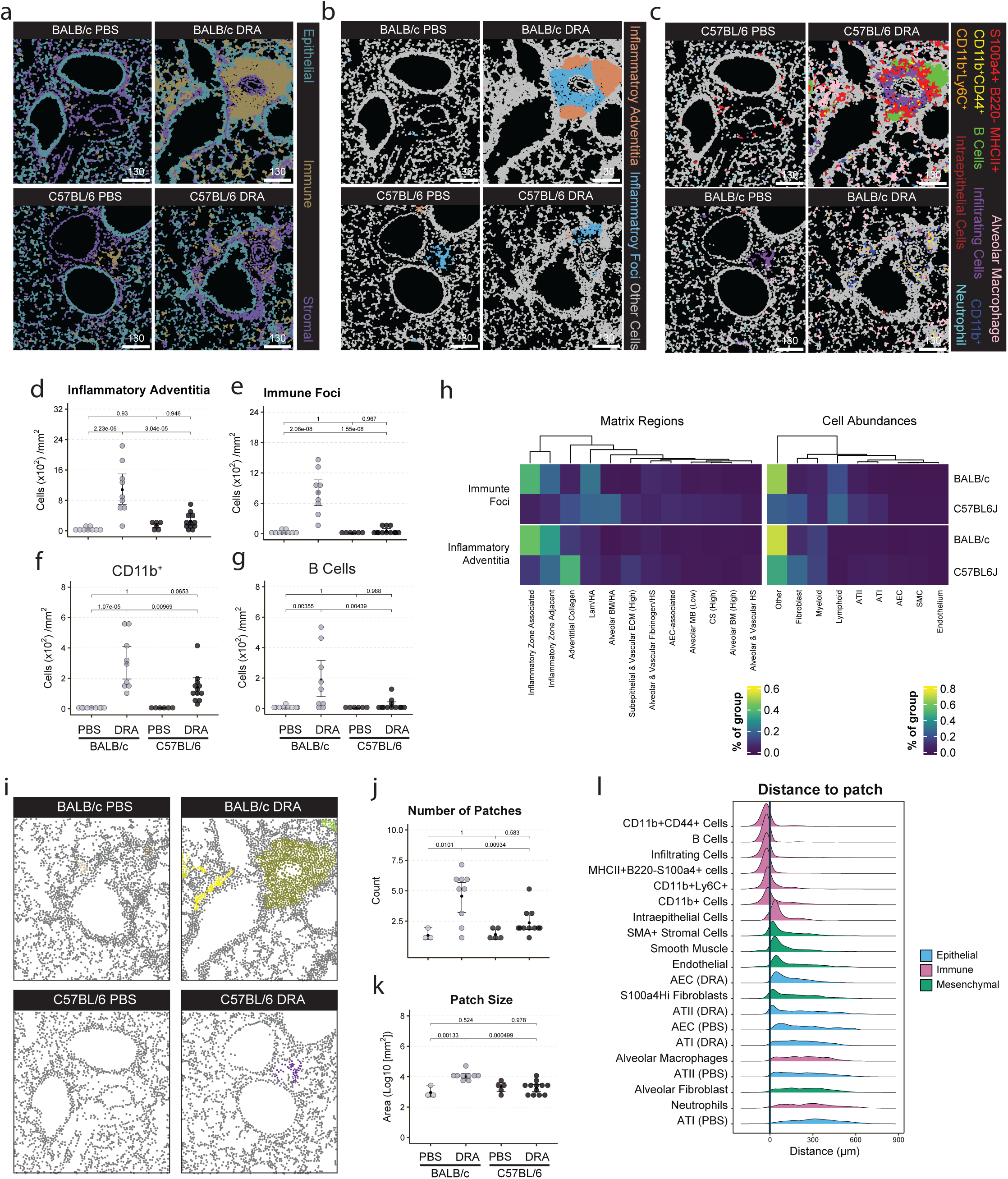
Allergic infiltrating immune cells localise in patches around the adventitial cuff. To classify the immune response to DRA challenge across the strains different immune cell populations were visualised. Projection of **a**) previously defined cell class, **b**) immune dominated lisaClust regions, and **c**) specific cell clusters on the deepcell masks. Quantification of the number of cells per mm^2^ within the **d**) inflammatory adventitia, **e**) immune foci lisaClust regions as well as **f**) CD11b^+^ and **g**) B Cells SNN clusters. **h**) To visualise the enrichment of specific matrix environments contingency tables were generated and plotted via heatmap. Left panel shows the percentage of cells in that region associated with the specific matrix environment. Right panel shows the percentage of cells in the region from previously defined cell classes. Scale represents the frequency of the region comprised of that matrix environment or cell type. To quantify if cells were accumulating together patchdetection() “patches” were calculated for the “immune” cell class. **i**) Representative images showing separate patches in unique colours for the four different groups. Quantification of **j**) the mean number and **k**) the mean area of the patches per ROI. **l**) The mean euclidian distance in μm of previously defined SNN cell clusters from their nearest patch. Vertical black line represents 0, the external border of the cell patch. Bars are coloured according to their previously defined cell class. **m**) Mean frequency of different cell types found across all patches across groups and **n**) mean frequency of specific SNN cell clusters within the immune cell class. For **d-g** and **j-k** data points represent individual ROIs with error bars showing the 95% confidence intervals around the mean (black filled circle). n = 2 - 4 animals per group corresponding to 6 - 12 ROIs per group. Statistical significance was calculated using a one-way ANOVA with Tukey post hoc comparison test. Scale bars equivalent to 130 μm.

To investigate the association of ECM components with these tissue regions contingency tables were generated for the defined matrix environments (**Fig 4 h**). The immune foci and inflammatory adventitia tissue regions were both enriched for the inflammatory zone adjacent and inflammatory zone associated matrix environments in BALB/c animals (**Fig 4 h**). These environments were generally sparse for ECM molecules (assayed in our panel) and only adjacent to type I and type III collagen (**Fig 3 e**). This aligned with an increased presence of “Lymphoid” and “other” cell types, mainly B cells and MHCII^+^B220^-^S100a4^+^ cells, in BALB/c mice (**Fig 4 h**). However, in C57BL/6 animals we found that these tissue regions were not only less prevalent but also associated with different matrix environments, particularly adventitial collagen and Lam/HA matrix environments (**Fig 4 h**). Taken together these results suggest that differences in matrix environments between BALB/c and C57BL/6 strains could be correlated with different immune cell compositions.

As BALB/c mice showed increased numbers of inflammatory cells which appeared to localise in distinct patches (**Fig 2 f** and **Fig 4 f and g**). We aimed to determine if these patches could be quantified, and their composition elucidated (**Fig 4 a**). Using the patchDetection() function^33^ we were able to identify contiguous patches of immune cells around the adventitial cuff in both PBS and DRA treated animals (**Fig 4 i**). Consistent with the lisaClust and cell population data (**Fig 4 d and e**), there were significantly greater numbers and size of patches in BALB/c DRA treated animals (**Fig 4 j and k**). Quantification of the relative distance of cell types to the detected patches demonstrated that immune cells (excluding intraepithelial cells, alveolar macrophages, and neutrophils) were found within patches, as evidenced by an average distance of <0μm from the patch boundary **(Fig 4 l**). Additionally, patches localised adjacent to smooth muscle, endothelium, and alveolar epithelial cells, consistent with their delimitation within the adventitial cuff (**Fig 4 i and l**). Despite being generated from immune cell clusters, epithelial and stromal cells were also contained within patches, suggesting a heterogenous tissue within the patch (**sup Fig 4 e and f**). While many inflammatory cells were found to reside within the adventitial cuff, some immune cells located specifically with the alveolar parenchyma, such as alveolar macrophages and neutrophils. Alveolar macrophages were increased in both strains in response to DRA treatment (**Fig 4 c, Fig 5 a**). Neutrophils were decreased following DRA challenge in BALB/c DRA relative to PBS treated animals and unchanged in C57BL/6 mice (**Fig 5 b**). Overall, these data suggest that inflammatory cells localise in distinct lung adventitial or alveolar tissue niches that correlate with specific matrix environments between strains.

**Figure 5:**
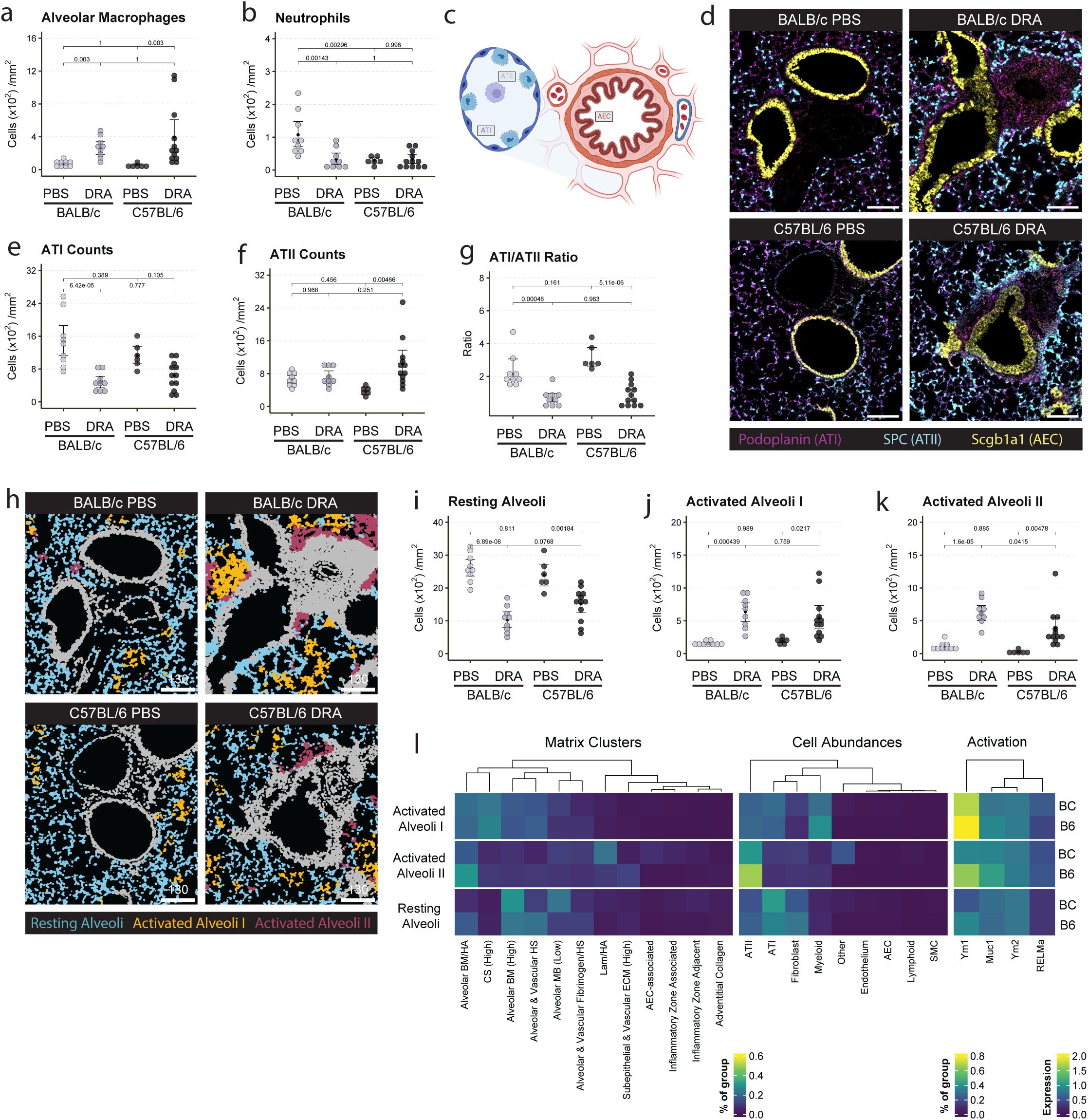
DRA induced AAI is associated with a loss of ATI and expansion of ATII. As we observed inflammatory cells also within the alveolar parenchyma we analysed the inflammatory changes within the epithelium during DRA challenge. Quantification of the mean cell number per mm^2^ of **a**) alveolar macrophages and **b**) neutrophils across ROIs. **c**) Schematic of the epithelial structure of the lung. **d**) Representative images showing staining of epithelial cell markers. Quantification of the mean cell number per mm^2^ of **e**) alveolar type I (ATI) **f**) alveolar type II (ATII) cells. **g**) Mean ratio of ATI to ATII cells per ROI. **h**) Images of the alveolar lisaClust regions projected onto representative ROIs. Quantification of the number of cells per mm^2^ in the **i**) resting alveoli, **j**) activated alveoli II, and **k**) activated alveoli I lisaClust regions. **l**) Visualisation of specific matrix environment enrichment. Contingency tables were generated and plotted via heatmap. Left panel shows the percentage of cells in that region associated with the specific matrix environment. Centre panel shows the percentage of cells in the region from previously defined cell classes. Both scales represent the frequency of the region comprised of that matrix environment or cell type. Right panel shows expression of specific activation markers, scale represents scaled and centred mean expression. For **a-b**, **e-g**, and **i-k** data points represent individual ROIs with error bars showing the 95% confidence intervals around the mean (black filled circle). n = 2 - 4 animals per group corresponding to 6 - 12 ROIs per group. Statistical significance was calculated using a one-way ANOVA with Tukey post hoc comparison test. Scale bars equivalent to 130 μm. Panel **c** generated using biorender.com.

### AAI is associated with a loss of ATI and expansion of ATII

The alveolar parenchyma has not often been considered a major site of remodelling during allergy. Three broad populations of lung epithelial cells; airway epithelial cells (AEC), alveolar type-1 cells (ATI), and alveolar type-2 cells (ATII) (**Fig 5 c**); were identified by expression of Uteroglobin (Scgb1a1) (AEC), Surfactant protein C (Spc) (ATII), and Podoplanin (Pdpn) (ATI) (**Fig 5 d, sup Fig 5 a-b).** ATI and ATII form the alveolar epithelium and are in contact with immune cells in the alveolar space. Whilst markers of epithelial activation (Ym1, Ym2, RELM, Muc1) separated epithelial cell type clusters by treatment (**Fig 2 d and e, sup 2a and sup Fig 5 c-f**), these clusters were merged for further analysis (**Fig 2 i**). ATI and ATII cells were significantly altered by DRA challenge, with reduced ATIs, albeit only significantly in BALB/c mice, (**Fig 5 e**) and increased ATII cells in C57BL/6 mice (**Fig 5 f**). ATI cell death followed by replacement via proliferation of ATII cells is typically associated with lung damage and repair^49–51^. To test for potential alveolar damage, the ratio of ATI and ATII cells was calculated. In both BALB/c and C57BL/6 animals the ATI:ATII ratio significantly decreased from approximately 3:1 in PBS controls to ∼1:1 in a DRA-treated animal (**Fig 5 g**), indicating a significant shift in the organisation of the alveoli during AAI.

As expected, ATI and ATII cells were the defining cells forming the alveolar parenchyma tissue regions **(Fig 2i-j).** Our analysis further separated the alveolar parenchyma into three lisaClust regions based on their exposure to PBS or DRA (**Fig 5 h**). These regions included the PBS dominated resting alveoli region and two activated alveoli which were more prevalent in DRA treated animals (**Fig 5 h-k**). To determine if there were specific ECM components associated with these alveolar tissue regions, we again generated contingency tables of the enriched ECM environments. Heparan sulphate, laminin γ 1, and type-IV collagen generally formed the basis of all the alveolar matrix environments (**Fig 3 e**). However, the resting alveoli were also characterised by the enrichments of numerous different matrix environments including alveolar basement membrane (High and Low) as well as alveolar & vascular heparan sulphate regions (**Fig 5 l**). These matrix environments showed differences between BALB/c and C57BL/6 PBS treated animals, aligning with our previous work highlighting strain-dependent differences in lung ECM molecules^21^. Both activated alveoli I and II regions showed disparate matrix environments correlating with the loss of ATI cells and alveolar fibroblasts (**Fig 5 e, l**, **and sup Fig 5 g and h**). Activated alveoli I were associated with CS (High) ECM environments, an increased prevalence of myeloid cells, and high expression of Ym1, likely coming from alveolar macrophages^52,53^ (**Fig 5 l and sup Fig 5 i**). Activated alveoli II regions were found adjacent to the inflammatory regions of the adventitial cuff (**Fig 5 h**). These regions were more prevalent in BALB/c mice (**Fig 5 j**), consistent with the increased immune infiltrates in this strain (**Fig 4 d and e**). These activated alveoli II regions were associated with increased proximity to Lam/HA regions in BALB/c mice and alveolar BM/HA regions in C57BL/6 mice (**Fig 5 l**). Overall, our data show distinct activation phenotypes of the alveolar epithelium can be defined during AAI using spatial analysis. Furthermore, these tissue regions are associated with specific matrix environments which may correlate with specific tissue functions.

### Airway epithelial cell activation is equivalent between strains

Airway epithelial cells (AEC) activation and subepithelial matrix deposition are key features of airway remodelling. To identify cells that may contribute to ECM changes during AAI, regions associated with the airways were analysed. The airways were divided into three distinct lisaClust regions: including resting and activated airways, as well as a population termed subepithelial cells which were localised basally underneath the airway epithelium (**Fig 6 a**). Additionally, lisaClust also isolated the associated blood vessel running parallel to these airways within the adventitial cuff (**Fig 6 a**). Following DRA administration, the number of cells within the resting airways decreased (**Fig 6 b and sup Fig 6 a**) and increased in the activated airways, as expected (**Fig 6 c and sup Fig 6 b**). The shift from resting to activated airway epithelium corresponded with increased Ym2 and RELMα expression in these cells (**Fig 6 d and sup Fig 6 and d**), molecules known to be associated with epithelial activation^21,54^. We observed no difference in the total number of AECs (**Fig 6 e**), nor any increase in cell diameter (**sup Fig 6 e**) or total airway circumference (**sup Fig 6 f**) between the strains before or after DRA challenge. However, when normalised to the diameter of the airways there was a significant increase in the cells/μm across the airways of C57BL/6 mice (**Fig 6 f**). This could be due to the expansion of AECs in response to DRA challenge in addition to a shrinkage of the airway due to excessive ECM deposition.

**Figure 6:**
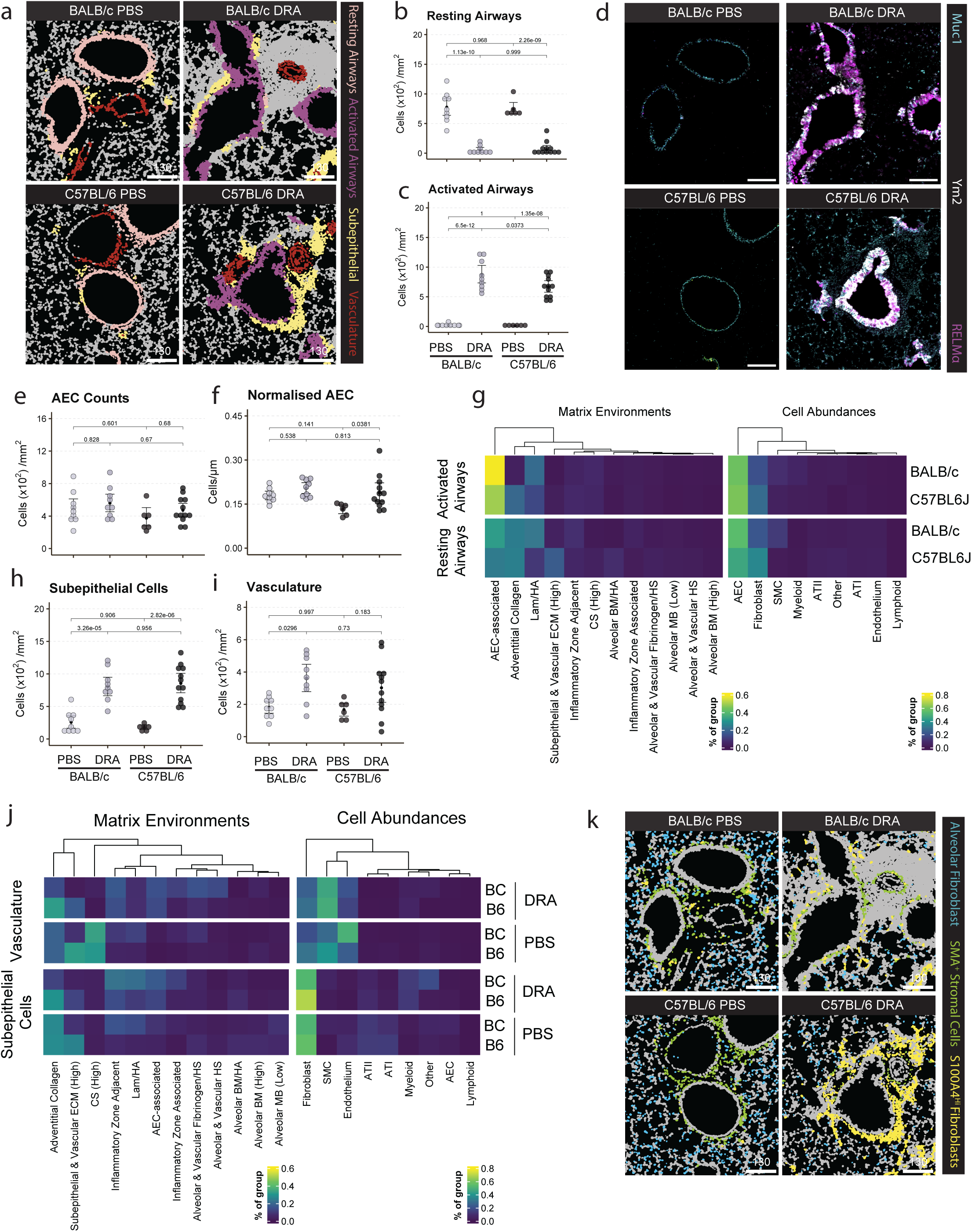
The subepithelial region is a site of immune-stromal interaction after DRA challenge. As the bronchial epithelium is a major site of remodelling during DRA induced allergic airway inflammation we sought to quantify the changes in this region. **a**) Projection of bronchial epithelium associated and vascular lisaClust regions onto deepcell masks. Quantification of the mean number of cells per mm^2^ in the **b**) resting airways and **c**) activated airways lisaClust regions. **d**) Representative images showing the expression of specific activation markers. Quantification of **e**) the total number of airway epithelial cells (AEC) and **f**) number of AECs normalised to the diameter of the airway in the respective ROI. **g**) To visualise the enrichment of specific matrix environments in the resting and activated airways lisaClust regions contingency tables were generated and plotted via heatmap. Left panel shows the percentage of cells in that region associated with the specific matrix environment. Right panel shows the percentage of cells in the region from previously defined cell classes. Representative images showing the localisation of airway and airway associated lisaClust regions in representative ROIs. Quantification of the mean cells per mm^2^ in the **h**) subepithelial and **i**) vasculature lisaClust regions. **j**) Visualisation of the enrichment of specific matrix environments in the subepithelial and vasculature lisaClust regions, contingency tables were generated and plotted via heatmap. Left panel shows the percentage of cells in that region associated with the specific matrix environment. Right panel shows the percentage of cells in the region from previously defined cell classes. **k**) Identified fibroblast populations projected onto deepcell masks of representative images. For **g** and **j** both scales represent the frequency of the region comprised of that matrix environment or cell type in the specified region across strains. For panels **b-c**, **e-f**, and **h-i** data points represent individual ROIs with error bars showing the 95% confidence intervals around the mean (black filled circle). n = 2 - 4 animals per group corresponding to 6 - 12 ROIs per group. Statistical significance was calculated using a one-way ANOVA with Tukey post hoc comparison test. Scale bars equivalent to 130 μm.

To determine if the alterations in the airways that we observed were correlated with changes in the surrounding ECM composition/organisation, we examined airway-associated matrix environments relative to cellular composition. Both resting and activated airways were linked with the AEC-associated matrix environment (**Fig 6 g**), characterised by proximity to hyaluronan, laminin γ1, and fibronectin. This signature was stronger in BALB/c mice and was further enriched in activated compared to resting airways (**Fig 3 e**). Notably the adventitial collagen matrix environment, enriched for type-I, III, and VI collagens, was downregulated in BALB/c activated airways but maintained in their equivalent C57BL/6 counterparts (**Fig 6 g**).

As there was a decrease in collagen containing matrix environments immediately proximate to the airways in BALB/c mice, but we had previously observed no difference in total subepithelial collagen^21^, we looked to characterise the matrix in the adjacent subepithelial region that was identified in our previous lisaClust analysis. There was a significant increase in the number of cells within this subepithelial region after DRA treatment in both strains (**Fig 6 h**). Increased cell numbers were also found in the vasculature region, although this increase was only significant in BALB/c animals (**Fig 6 i**). We then split the vascular and subepithelial regions by treatment to interrogate differences with and without DRA challenge. The vasculature region was comprised of smooth muscle cells (SMC) and endothelial cells alongside a smaller number of fibroblasts. In PBS treated animals this vasculature region was dominated by a CS high matrix environment, but this was almost completely lost after DRA treatment, suggesting a downregulation of vascular CS expression during allergy (**Fig 6 j**). This was accompanied by an increase in the prevalence of SMC, suggesting a remodelling of the vessels during DRA challenge.

In contrast, the subepithelial region had a similar matrix environment across PBS and DRA treated animals. This region was enriched for adventitial collagen and subepithelial & vascular ECM regions (**Fig 6 j**) comprising both type I and III collagens, two of the major isoforms deposited under the airways in this model^21^ (**Fig 3 e**). The cellular composition of the subepithelial region was primarily fibroblasts (**Fig 6 j**) matching with the localisation of two of the three stromal populations identified, namely SMA^+^ stromal cells and S100a4^+^ fibroblasts, compared to alveolar fibroblasts localised found in the alveolar parenchyma (**Fig 6 k** and **Fig 2**). In addition, one of the few populations that appeared enriched within the subepithelial region during DRA challenge in both strains were myeloid cells (**Fig 6 j**) suggesting these cells may be infiltrating into this region of active ECM remodelling during AAI.

### CD11b^+^ immune cells localise close to S100a4^+^ subepithelial fibroblasts during AAI

As we had identified CD11b^+^ cells infiltrating into the subepithelial region we aimed to determine if these cells were interacting with the resident stromal populations. Within the subepithelial region SMA^+^ stromal cells and S100a4^+^ fibroblasts localised close to collagen-I and III, corroborating analysis of the matrix environment contingency plot (**Fig 7 a and b**). SMA^+^ stromal cells were present in PBS treated animals and did not increase following DRA administration (**Fig 7 c**). However, S100a4^+^ fibroblasts were significantly expanded in both allergic BALB/c and C57BL/6 animals (**Fig 7 d**). Additionally, in DRA animals we were able to identify a population of CD11b^+^ cells near to both stromal populations in the subepithelial region of DRA treated mice (**Fig 7 e**).

**Figure 7:**
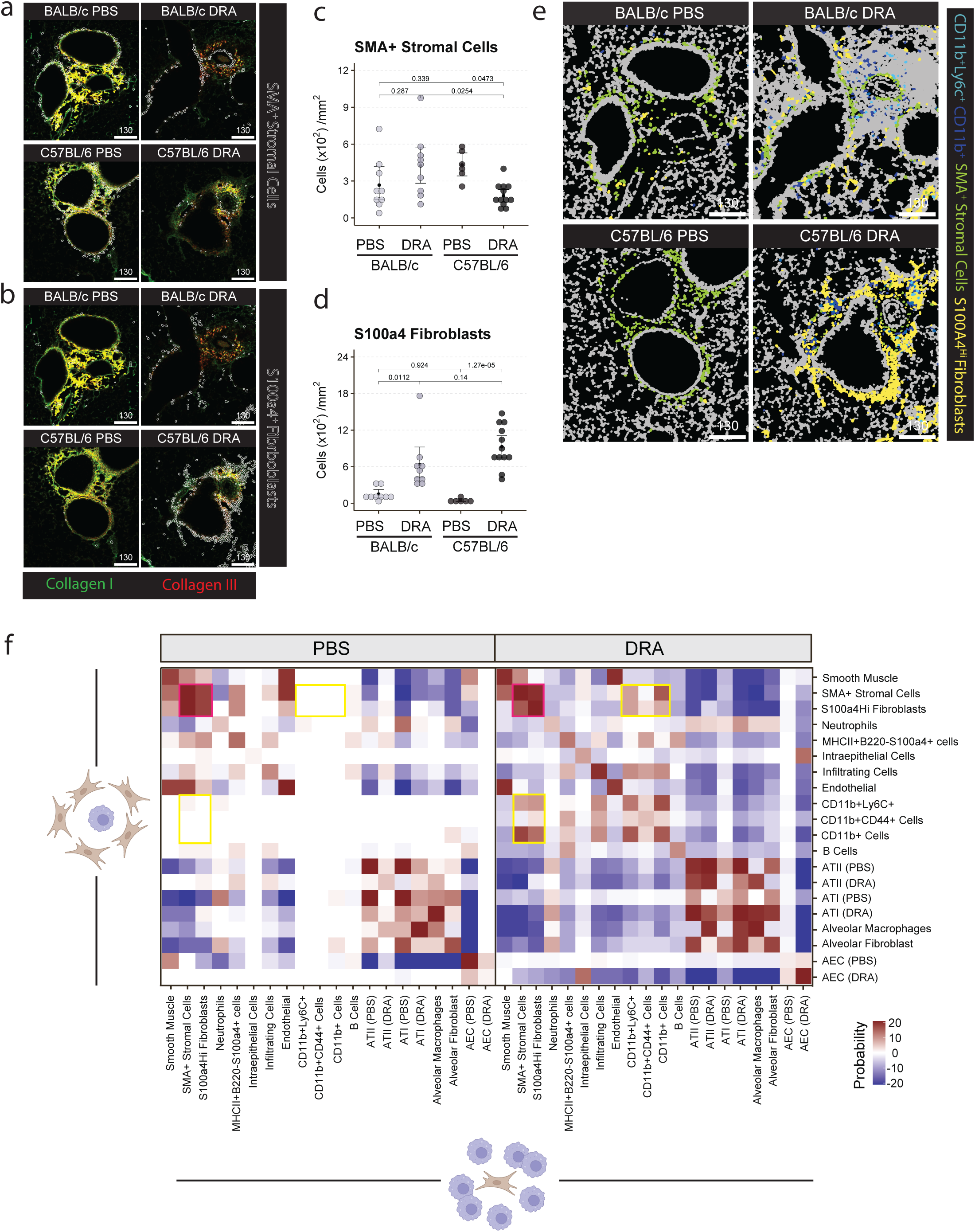
S100a4+ subepithelial fibroblasts localise with CD11b+ immune cells during AAI. As there was an enrichment of stromal cells in the area of remodelling we sought to further interrogate the interaction of these cells with the ECM. Representative images showing the localisation of **a**) SMA^+^ stromal cells and **b**) S100a4^+^ fibroblasts in relation to type-I and type-III collagen. Quantification of the mean number per mm^2^ of **c**) SMA^+^ stromal cells and **d**) S100a4^+^ fibroblasts per ROI. **e**) Representative images showing the projection of key stromal and CD11b^+^ cell populations on deepcell generated masks. **f**) Heatmap showing the probability that distinct cell populations are found in close proximity more than random chance in PBS and DRA animals, as calculated using neighbourhood analysis^95^. For panels **c-d** data points represent individual ROIs with error bars showing the 95% confidence intervals around the mean (black filled circle). n = 2 - 4 animals per group corresponding to 6 - 12 ROIs per group. Statistical significance was calculated using a one-way ANOVA with Tukey post hoc comparison test. Image scale bars are equivalent to 130 μm.

To clarify whether cells within the subepithelial region were in close association and likely to be interacting, we utilised neighbourhood analysis^27^. Cell neighbours were determined using Euclidian distance between cell masks to define which cells were close to each other. Then the proportion of certain cell types was compared to a randomly generated distribution to quantify if specific populations were over or under-represented. Analysis demonstrated an overrepresentation of interactions between CD11b^+^ and CD11b^+^Ly6C^+^ cells with SMA^+^ stromal cells and S100a4^+^ fibroblasts only in the DRA treated samples (*yellow box*) (**Fig 7 f**). SMA^+^ stromal cells and S100a4^+^ fibroblasts were found to be associated in both PBS and DRA challenged animals (*magenta box*) (**Fig 7 f**) suggesting that this interaction may also be important also in the steady state.

Together, these data suggests that a subset of CD11b^+^ immune cells are accumulating in the subepithelial niche during allergen challenge. Here, these cells can interact with SMA^+^ stromal cells and correlate with an expanded S100a4^+^ fibroblast population. As ECM remodelling occurs predominantly in the subepithelial region, these cell interactions may form an important network for regulating ECM deposition during allergic pathology.

## Discussion

We have characterised three distinct regions of the lung that are altered during allergic airway inflammation. By combining cellular profiling and extracellular matrix (ECM) phenotyping we identify distinct matrix environments that are associated with areas of tissue remodelling during allergic disease. Studies in murine models of allergic airway inflammation (AAI) have traditionally focused on small and medium airways as the sites of remodelling. However, during DRA induced AAI we observed numerous changes in the alveolar tissue structure, including altered ATI/ATII balance, alveolar cell activation, and the expansion of alveolar macrophages. Tissue resident alveolar macrophages are generally tissue protective^55^ and have been shown to prevent neutrophil-mediated tissue damage^56^. During allergen sensitisation, these resident cells are replaced^57^, mainly through recruitment and differentiation of circulating monocytes but also proliferation of these populations^58^. This expansion of recruited macrophages then corresponds to alterations in morphology and surfactant dysfunction within the alveolar epithelium^57^. These changes in alveolar epithelial cells are then key for regulating subsequent immune responses^57,59,60^. The exact roles that the alveolar epithelium plays in the pathophysiology of AAI and its impact on tissue function are poorly understood.

Distinct parenchymal ECM environments were observed in the lungs of DRA-challenged mice. This aligns with human studies showing matrix turnover in the alveolar spaces differs between controlled and uncontrolled asthma^61^. Specifically, we observed increased hyaluronan and chondroitin sulphate in the alveolar compartment. These are ECM components important for regulating cell infiltration into the tissue^62–64^, hence these molecules are potentially important in regulating the immune tissue niche and controlling alveolar macrophage and neutrophil localisation. Fitting with this, chondroitinase treatment of bleomycin-induced fibrotic murine lungs reduced airway macrophages^65^. Similarly, studies of fibrotic remodelling in hamsters, demonstrate a switch from heparan sulphate dominated to chondroitin sulphate dominated matrices^66^, changes likely driven by alterations in fibroblasts and epithelial cells. Alveolar fibroblasts are known to take on different phenotypic states during inflammatory and fibrotic responses^67^. In our study, alveolar fibroblasts decreased upon DRA challenge. However, rather than dying, these cells are likely becoming activated/differentiating leading to incorporation into other cell clusters within our dataset. Identifying specific markers will aid a better understanding of how alveolar fibroblast lineage progression may influence response in health versus disease.

In asthma and models of AAI, there is no doubt that the airways are a key site of pathological changes which have many downstream consequences for lung function. We have identified a region of subepithelial tissue, adjacent to the smooth muscle layer and expanded during DRA challenge in both strains. Alongside enrichment for fibroblasts, this subepithelial region was also associated with the infiltration of CD11b^+^ immune cells. Although we were unable to subdivide this cell cluster further in our dataset, eosinophils, interstitial macrophages, and monocytes are all possibly present. Fibroblast-immune cell networks have been described previously, albeit largely based on *in vitro* assays. Eosinophils and fibroblasts have been shown to cross-regulate one another via the production of several soluble mediators^68^, with eosinophils isolated from people with asthma inducing a distinct fibroblast phenotype associated with ECM remodelling^69^. Interstitial macrophages, which express high levels of CD11b, are a heterogenous cell population and their true diversity is only recently being explored^70–72^. Interstitial macrophages increase in the lungs of mice during fibrosis^73^, and DRA-induced AAI^21^, but their exact influence on disease progression are unclear. CpG induced expansion of IL-10-producing interstitial macrophages was protective during HDM-induced AAI^74^. However, depletion of interstitial macrophages in radiation-induced lung fibrosis has protective effects^75^. Together this suggests that their roles may be more complicated and may be segregated to different functional cell subtypes. It has been shown that macrophages can take and provide cues to fibroblasts^76^, in line with observations in our study which show CD11b^+^ and CD11b^+^Ly6c^+^ cells were closely interacting with expanded S100a4^+^ fibroblasts and SMA^+^ stromal cells within the subepithelial tissue region. These interactions may be particularly important considering fibroblast S100a4 expression increases over time with inflammation and fibrosis^67^. Recent data from Yu *et al.* identified a fibroblast population associated with numerous contractile genes interacting with macrophages by a Cxcl12/Cxcr4 pathway^77^. Within our data the subepithelial tissue ECM environment showed an enrichment for type VI collagen, amongst other ECM molecules. Recent studies demonstrate that during pulmonary fibrosis, macrophages within the interstitial space influence fibroblast function through secretion of type VI collagen^78^. Furthermore, type VI collagen can then prevent fibroblast apoptosis^79^ and is essential for fibroblast migration^80^. Together, our spatial studies support the idea of bidirectional communication between fibroblasts and macrophages within the subepithelial region. Therefore, it is interesting to propose that one way in which fibroblasts and macrophages may communicate is via alterations to their surrounding ECM. Further investigation into how ECM environments shape and regulates cell-cell interactions, cellular activation and spatial localisation within regional tissue architecture may provide avenues to restore lung homeostasis following pathology.

Our study begins to computationally explore the spatial relationship between cells and the ECM during chronic allergic pathology. Integration of cellular signals with spatial ECM analysis in further studies will be an important step given the key role the ECM plays in shaping cellular responses^9,81^. In addition many key pulmonary immune and stromal populations are beginning to be described by single cell RNA sequencing^82,83^, but do not yet have good markers for antibody-based imaging approaches. Alongside this cellular RNA profiles of ECM-related genes are often poor correlators with the surrounding bioactive ECM molecules. Therefore, interrogation of the ECM network is only truly accessible by direct analysis of the molecules themselves. Work is being done on integrating pathway based analysis of ECM regulation using single cell transcriptomic data^84^ however, molecular and spatial analysis will always be required to confirm these findings. Regardless, the more detail from the *in vivo* setting we are able to recapitulate in our analysis, the greater our understanding of how cell-cell and cell-ECM networks support health versus disease. Developing additional computational image analysis processes for studying these cellular and ECM datasets, will only aid in identifying new pathways that regulate inflammation and lung function during disease.

## Materials and Methods

### Animals

BALB/c^olaHsd^ and C57BL/6J^olaHsd^ mice were obtained from a commercial supplier (Envigo, Hillcrest, UK). All experimental animals were female and 8 weeks of age at the start of the experiment. Mice were housed in individually ventilated cages in groups of five animals in specific pathogen-free facilities at biological services facility the University of Manchester. Mice were randomly assigned to a treatment group. Sample size was calculated based on pilot experiments (N=3 per group), based on the number of animals needed for detection of a 25% change in Masson’s trichrome-positive area around the airway in PBS *versus* allergic mice (Power = 0.8, P-value <0.05). All animal experiments were performed in accordance with the UK Animals (Scientific Procedures) Act of 1986 under a Project License (70/8548) granted by the UK Home Office, approved by the University of Manchester Animal Welfare and Ethical Review Body. Animals were euthanised by asphyxiation in rising concentrations of carbon dioxide and confirmation by cessation of circulation.

### Allergic airway inflammation model

A model of allergic airway inflammation was induced in animals as previously described^20,21^. Allergen cocktail containing 5µg house dust mite (*Dermatophagoides pteronyssinus,* 5450 EU, 69.23mg per vial), 50µg ragweed (*Ambrosia artemisiifolia*) and 5µg *Aspergillus fumigatus* extracts (Greer Laboratories, Lenoir, NC, USA) was freshly prepared prior to installation. Mice were administered with either 20µL of DRA cocktail or PBS via intranasal instillation. This occurred twice weekly for up to 8 weeks following anesthetisation by isoflurane inhalation. Mice were then rested for 5 days prior to collecting lung tissue.

### Sample preparation

The lone lobe was fixed overnight in 10% neutral buffered formalin (NBF) before being stored in 70% ethanol. Tissues were then processed and embedded in paraffin wax using standard protocols. Tissue sections (5µm) were prepared using a microtome onto glass slides (Superfrost Plus adhesion). Tissue sections were cut immediately prior to staining with antibodies for imaging mass cytometry (IMC).

### IMC antibody conjugation

Purified, carrier-free antibodies (see Table. 1) were conjugated to selected metal isotopes using the MaxPar antibody conjugation kit and following manufacturer’s instructions (Standard BioTools). Briefly, Ln metals were loaded onto MaxPar polymer and incubating for

60 minutes at room temperature (RT), with regular agitation. Antibodies were partially reduced and coupled with metal-loaded polymer before being washed and concentrated to a volume of approximately 20 µL using a 50 kDa molecular weight cut off filter column. Final concentrations of conjugated antibody solutions were measured and antibodies stored in stabilisation buffer (Candor) at 4°C.

**Table 1:**
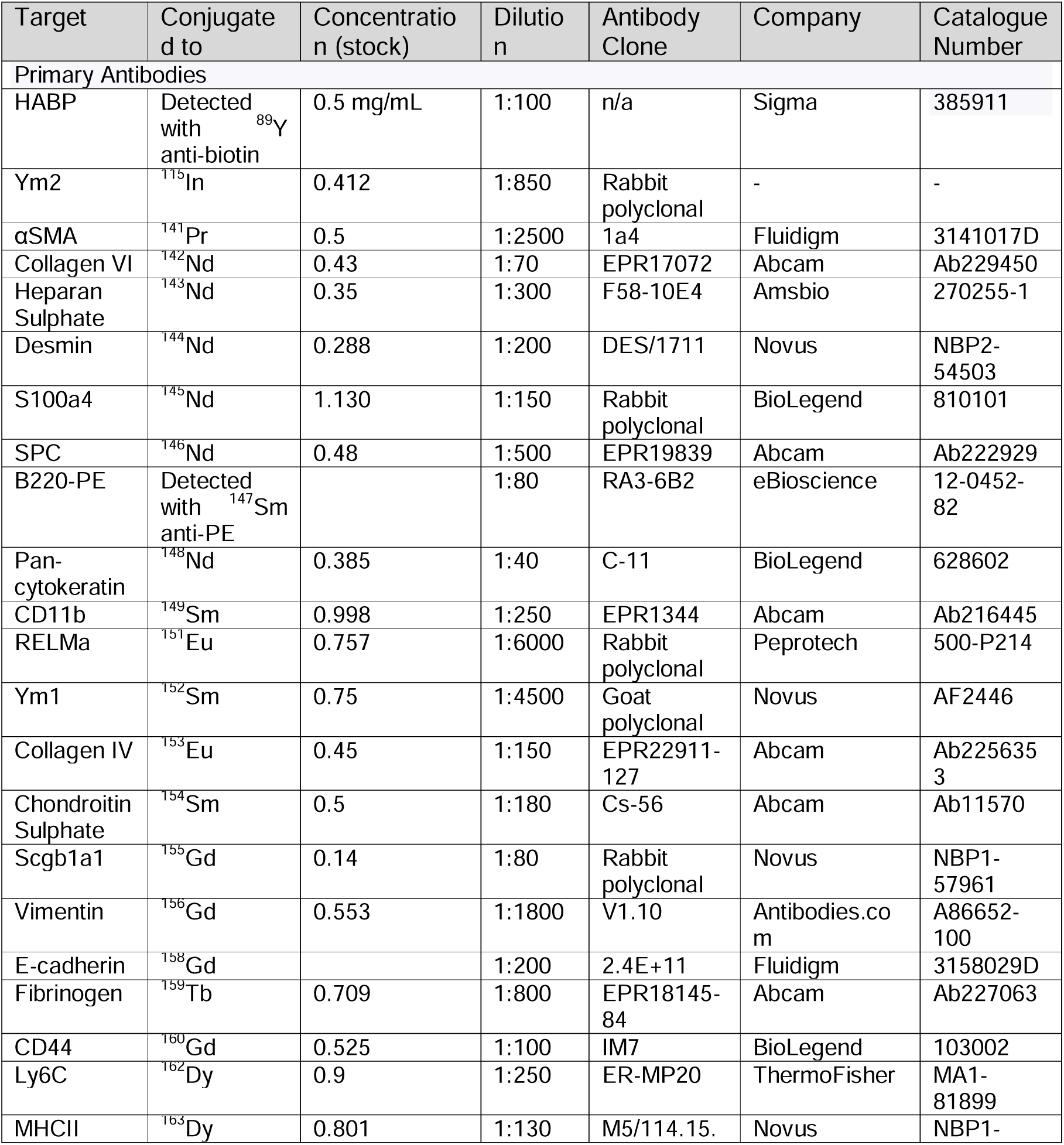
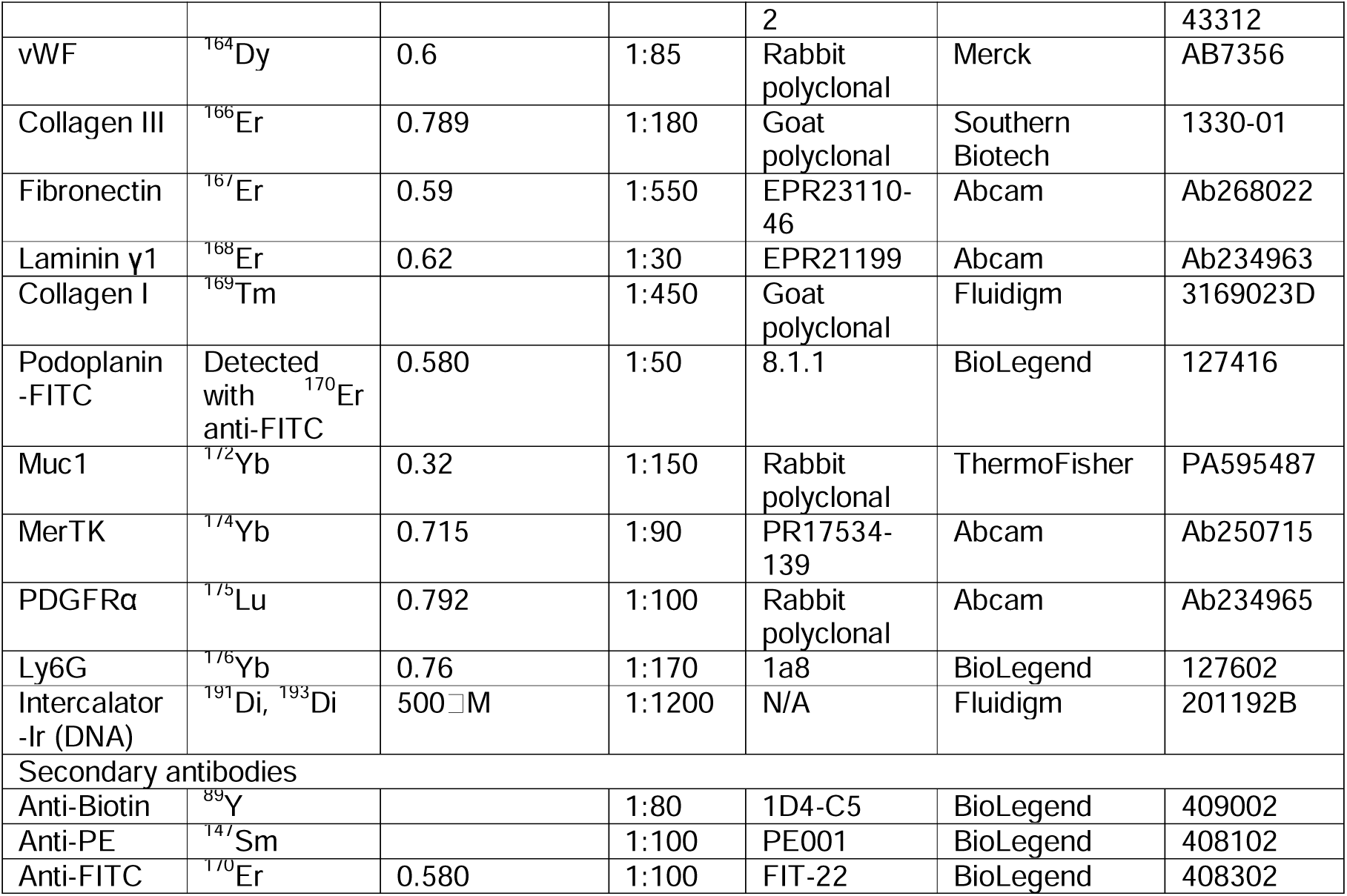
Antibodies for tissue staining.

### IMC staining

Tissue sections were dewaxed through xylene and an ethanol gradient using standard protocols. Antigen retrieval was performed on dewaxed sections using 10 mM Tris-HCl, pH 9.0, 1 mM ethylenediaminetetraacetic acid (EDTA) + 0.02% Tween-20 for 20 minutes by heating in a microwave. Slides were then permeabilised in 0.2% Triton-X100 in Phosphate Buffered Saline, pH 7.4 (PBS) for 20 minutes at RT prior to staining. Non-specific protein was blocked with 3% Bovine Serum Albumin (BSA) in PBS for 1 hour at RT in a hydration chamber. For staining, antibody cocktails (see Table. 1) were prepared in PBS with 0.5% BSA. All conjugated antibodies were centrifuged (10,000 *g* for 2 mins) immediately prior to use to minimise antibody aggregates. Sections were incubated with in primary antibody cocktail overnight at 4°C in a hydration chamber. Slides were then washed in PBS with 0.2% Triton-X100 (8 minutes), followed by PBS alone (8 minutes). Secondary antibodies (see Table. 1) were diluted in PBS with 0.5% BSA as before and added to tissue sections for 1 hour at RT. Slides were then washed as before and incubated with Iridium Intercalator in PBS (1:1200) for 30 minutes at RT. Stained slides were washed in dH_2_O for 2 minutes with gentle agitation and then air-dried for at least 20 minutes at RT and stored at 4°C until acquisition.

### Image collection

IMC data was acquired on the Hyperion® Imaging System (Standard BioTools). The instrument was calibrated using a tuning slide to optimise the sensitivity of the detection range with a signal for 175Lu of at least 1500 mean duals used. Specific 660 x 690μm regions of interest (ROI) were laser-ablated in 1μm^2^ pixels before being passed through a plasma source and metal-conjugated antibodies detected with time-of-flight mass spectrometry. Antibody staining was reviewed with histoCAT++ and data extracted from mcd files.

### Deepthresh matrix thresholding

All regions of interest (ROIs) were first enhanced using IMCDenoise^89^. Six ROIs were manually thresholded for use in a semi-automated thresholding approach we refer to as DeepThresh. Four images were used for training, one for validation and one for testing. DeepThresh was built using PyTorch^90^ (version 2.3) and Pytorch Segmentation Models^91^. The architecture mimics that of U-Net^92^ with a ResNet152 encoder^93^. Adam^94^ optimizer was applied with a learning rate of 0.0001. All models were trained for 50 epochs. In relation to training, a separate DeepThresh model was trained for each channel. ROI’s were channel-wise min-max normalised. Each image was zero padded around the edges to enable splitting into overlapping 64×64 patches with a stride of 32. In addition, data augmentation was applied by rotating the patches by 90° and 180°. In terms of inference, the images were zero padded, split into 64×64 non-overlapping patches, normalised and the binary mask predicted from the trained channel specific models. Subsequently the patches were reconstructed and the zero-padding removed.

### Matrix distance calculation

To accurately analyse ECM molecules, we developed a pipeline using binary thresholding of matrix staining, followed by distance calculation between positive ECM staining and previously generated cell masks (**sup. Fig 3 a**). Firstly, processed stacked images were exported from the IMCdenoise^89^ pipeline (**sup. Fig 3 a i**). Manual thresholding was applied to a subset of images to generate binary images of positive ECM marker staining (**sup. Fig 3 a ii**). The binary images were then used as training data in a semi-automated thresholding approach referred to as DeepThresh (**sup. Fig 3 a iii & sup Fig 3 b**). DeepThresh was then applied to the entire dataset which allowed thresholding of ECM channels whilst accounting for variances in brightness and background between ROIs (**sup. Fig 3 a iv**). Once thresholding had been applied, skimage^44^ was used to calculate distance maps from binary images (**sup. Fig 3 a v**). Distance maps represent the value of each pixel as its distance from the closest positive pixel in the binary thresholded image. Distance maps were appended to z stack images prior to importing into the Steinbock pipeline enabling ECM-cell data integration into the standard pipeline (**sup. Fig 3 a vi**). Quantification of average values across deepcell generated cell masks, as implemented in the steinbock package, then gave the average distance of each cell from specific ECM components (**sup. Fig 3 a vii**).

### IMC image preparation and analysis

Following data extraction, cell mask generation, and cell marker quantification IMC data was imported into R using the imcRtools package. Batch correction was performed using the R implementation of Harmony, providing the slide number as a variable, to remove variance between different slides when imaged at different times. Following batch correction, dimensionality reduction and clustering were performed using principle component analysis (PCA)^85,86^, multidimensional scaling (MDS)^87^, uniform manifold approximation projection (UMAP)^88^, and single nearest neighbours (SNN)^45^ respectively.

Sup. Fig 2 Additional cell phenotyping information. **a**) Heatmap of a subset of 20,000 cells showing the arcsin() transformed mean expression per cell of the different markers. **b**) Multiple dimension scaling (MDS) of cell marker expression. **c**) MDS scaling using the proportions of different cell clusters per sample. **d**) lisaClust regions across all ROIs. **e**) Quantification of mean cell diameter per ROI. For **e** data points represent individual ROIs with error bars showing the 95% confidence intervals around the mean (black filled circle). n = 2 - 4 animals per group corresponding to 6 - 12 ROIs per group. Statistical significance was calculated using a one-way ANOVA with Tukey post hoc comparison test.

Sup. Fig 3: To quantify matrix distances, we developed a new pipeline for thresholding matrix staining.

**a)** Schematic showing the DeepThresh pipeline for matrix distance quantification. **i**) Raw stacked tiff data was exported from IMCdenoise pipeline and split into two groups. **ii**) One group consisting of a representative sample from each biological group is manually thresholded to generate binary image showing positive staining as 1 (foreground) and negative staining as 0 (background). **iii**) A random forest classifier was trained using this data to identify positive regions. **iv**) The trained classifier was then applied to the full dataset to generate thresholded data. **v**) Thresholded images were used to calculate matrix pixel distances using the scikit image package and these images were appended to the original stacked tiff files. **vi**) Masks were generated with deepcell using the IMCdenoise data and were then used to quantify the expression of cell markers and distance values per cell mask. Which was imported into the normal steinbock pipeline for further analysis. **b**) Description of the methodology for thresholding images using DeepThresh. All regions of interest (ROIs) were first enhanced using IMCDenoise^89^. Six ROIs were manually thresholded for use in a semi-automated thresholding approach we refer to as DeepThresh. Four images were used for training, one for validation and one for testing. DeepThresh was built using PyTorch2 (version 2.3) and Pytorch Segmentation Models3. The architecture mimics that of U-Net4 with a ResNet152 encoder5. Adam6 optimizer was applied with a learning rate of 0.0001. All models were trained for 50 epochs. **c**) Heatmap showing the matrix distances across classified matrix environments for all ECM markers evaluated. Scale shows reciprocal transformed.

Sup. Fig 4 Additional quantification of cell populations. Quantification of mean cells per mm^2^ of **a**) CD11b^+^CD44^+^ cells, **b**) Infiltrating cells, **c**) CD11b^+^Ly6C^+^ cells, **d**) MHCII^+^B220^-^S100a4^+^ cells per ROI. For **a-d** data points represent individual ROIs with error bars showing the 95% confidence intervals around the mean (black filled circle). n = 2 - 4 animals per group corresponding to 6 - 12 ROIs per group. Statistical significance was calculated using a one-way ANOVA with Tukey post hoc comparison test.

Sup Fig 5 Additional phenotyping of the allergic epithelium. Violin plot showing **a**) SPC expression and **b)** Podoplanin expression in epithelial subsets. Quantification of **c**) resting ATI, **d**) activated ATI, **e**) resting ATII, **f**) activated ATII epithelial clusters. **g**) Representative images showing localisation of alveolar fibroblasts (white outlines) and type-I and type-III collagen. **h**) Quantification of the mean number of alveolar fibroblasts per ROI. **i**) Enrichment of alveolar macrophages across epithelial lisaClust region in different treatment groups. For panels **a-b** violin plots represent the mean expression calculated across the cell mask. For **c-f** and **h** data points represent individual ROIs with error bars showing the 95% confidence intervals around the mean (black filled circle). n = 2 - 4 animals per group corresponding to 6 - 12 ROIs per group. Statistical significance was calculated using a one-way ANOVA with Tukey post hoc comparison test. Scale bars equivalent to 130 μm.

Sup. Fig 6 Additional quantification of the airways. Quantification of the mean cells per mm^2^ of **a**) Resting airway epithelial cells (AEC) and **b**) activated AEC per ROI. Violin plots showing the mean expression per cell of canonical epithelial activation markers **c**) Ym2, and **d**) RELMα. Quantification of the mean diameter **e**) and total circumference **f**) of AEC per ROI. For panels **a-c** data points represent individual ROIs with error bars showing the 95% confidence intervals around the mean (black filled circle). For panels **d-f** violin plots represent the mean expression calculated across the cell mask. n = 2 - 4 animals per group corresponding to 6 - 12 ROIs per group. Statistical significance was calculated using a one-way ANOVA with Tukey post hoc comparison test.

Sup Fig 7 Additional analysis of the smooth muscle layer. Quantification of **a**) the mean number of smooth muscle cells per ROI and **b**) the mean diameter of smooth muscle cells per ROI. For panels **a-b** data points represent individual ROIs with error bars showing the 95% confidence intervals around the mean (black filled circle). For panels **d-f** violin plots represent the mean expression calculated across the cell mask. n = 2 - 4 animals per group corresponding to 6 - 12 ROIs per group. Statistical significance was calculated using a one-way ANOVA with Tukey post hoc comparison test.

## Acknowledgements

The authors thank Jennifer Baron and Gareth Howell, and the Flow Cytometry core facility at the University of Manchester for their Hyperion imaging technical support service as well as Matthew Burgess and Morgan Bryant for critically reading the manuscript. Thanks should also go to Stella Pearson for laboratory technical support. For the purpose of open access, the authors have applied a Creative Commons Attribution (CC BY) licence to any author accepted manuscript version arising from this submission.

## Author contribution

Conceptualisation: JEP, RJD, and TES

Methodology: JEP, MF, and MG

Software: JEP, MF, and MG

Formal analysis: JEP and MG

Investigation: JEP and TES

Writing - Original Draft: JEP

Writing - Review & Editing: JEP, TES, RJD, HET, MG, MR

Visualization: JEP and MG

Supervision: TES, JEA, and MR

Funding acquisition: TES, JEA, and MR

