## Supplemental Figures for "Extracellular matrix phenotyping by imaging mass cytometry defines distinct cellular matrix environments associated with allergic airway inflammation"

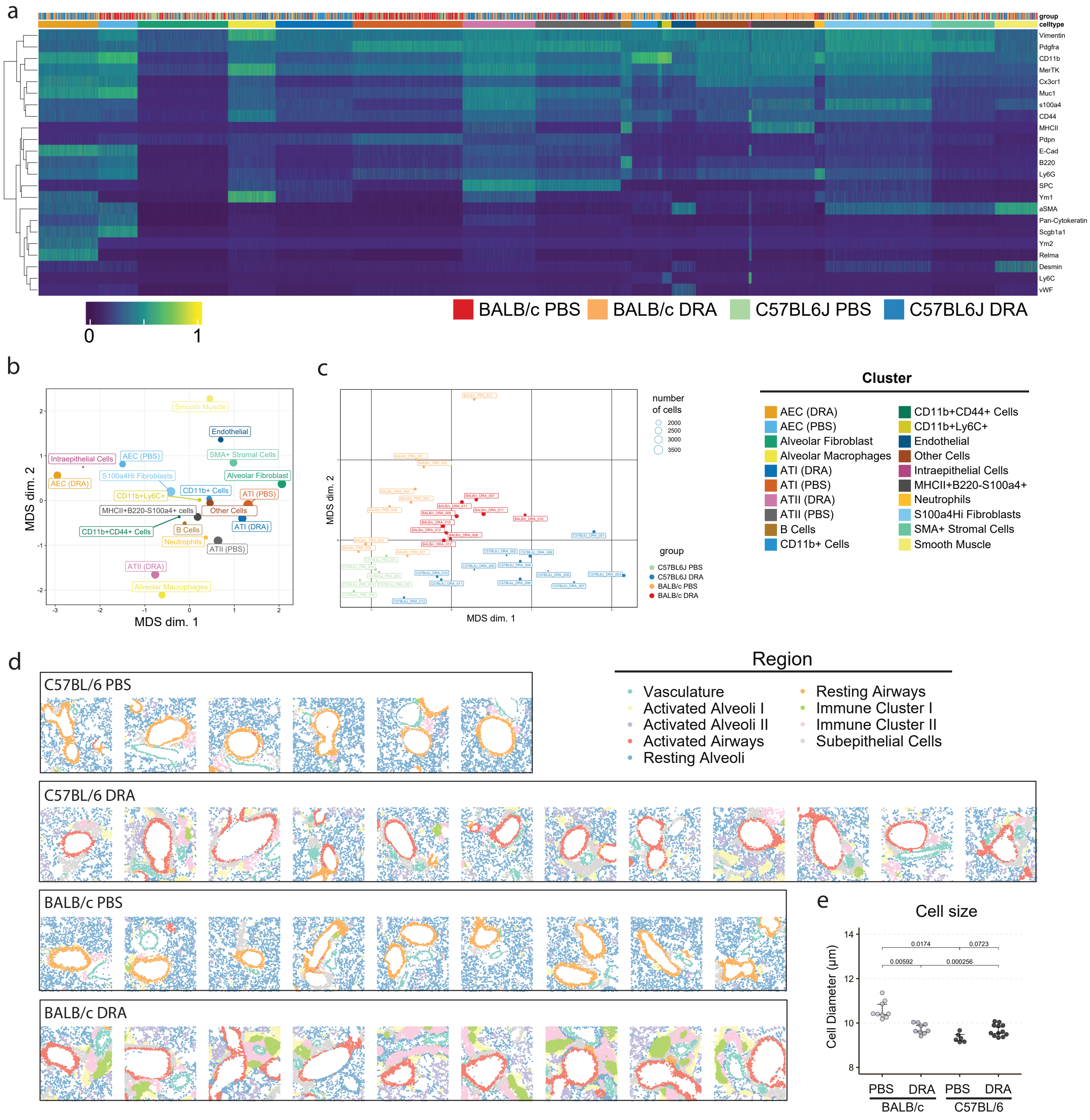


Supplementary Figure 2

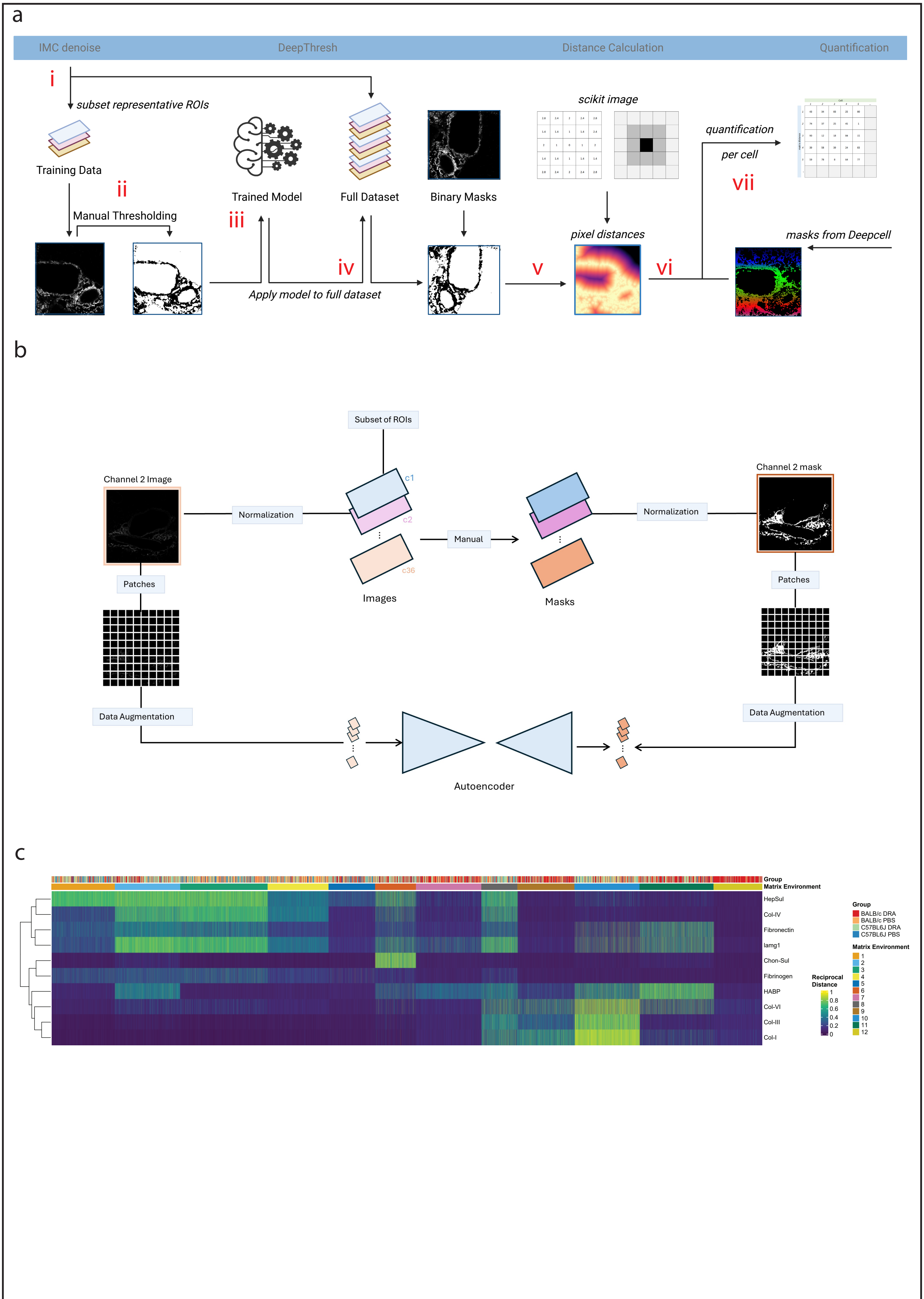


Supplementary Figure 3

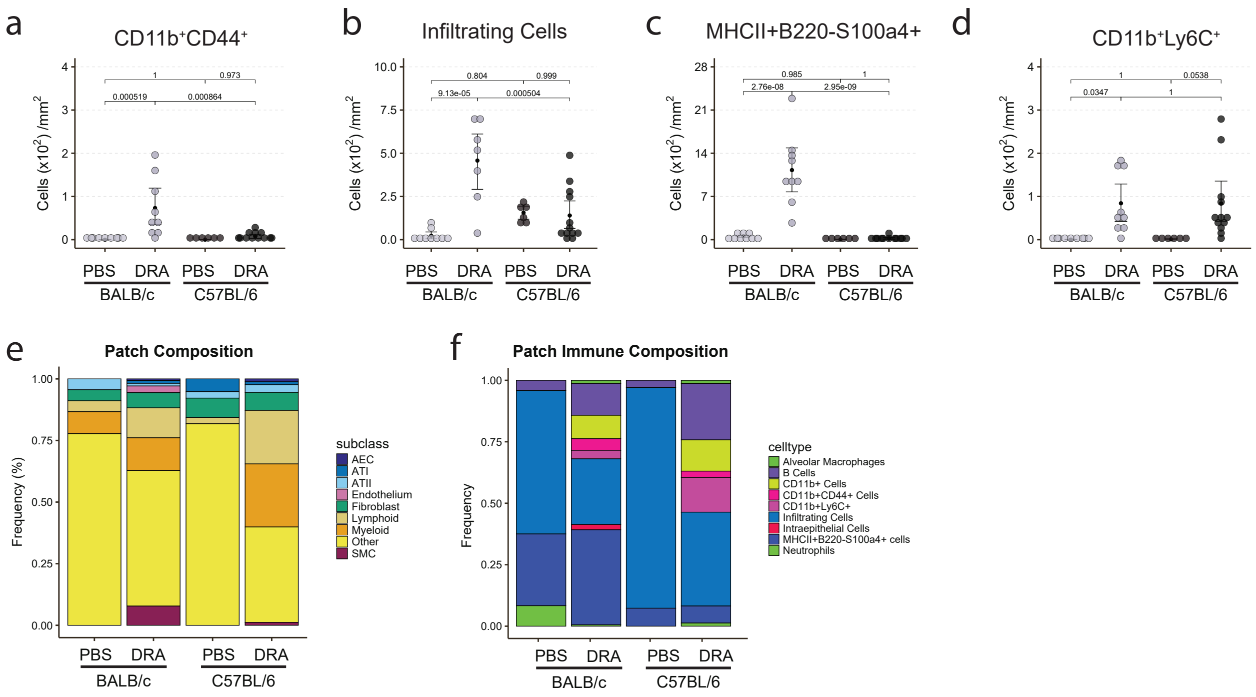


Supplementary Figure 4

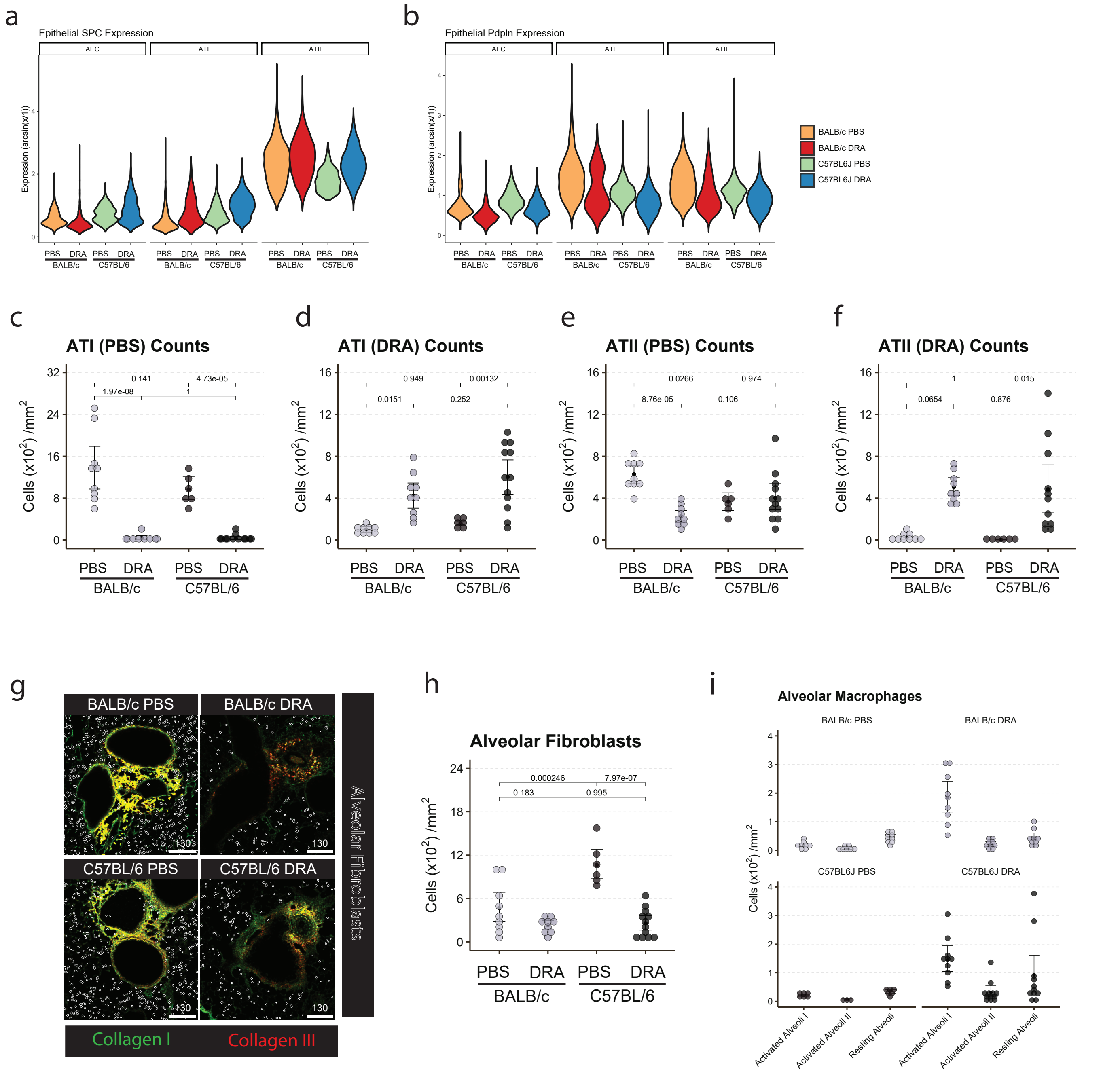


Supplementary Figure 5

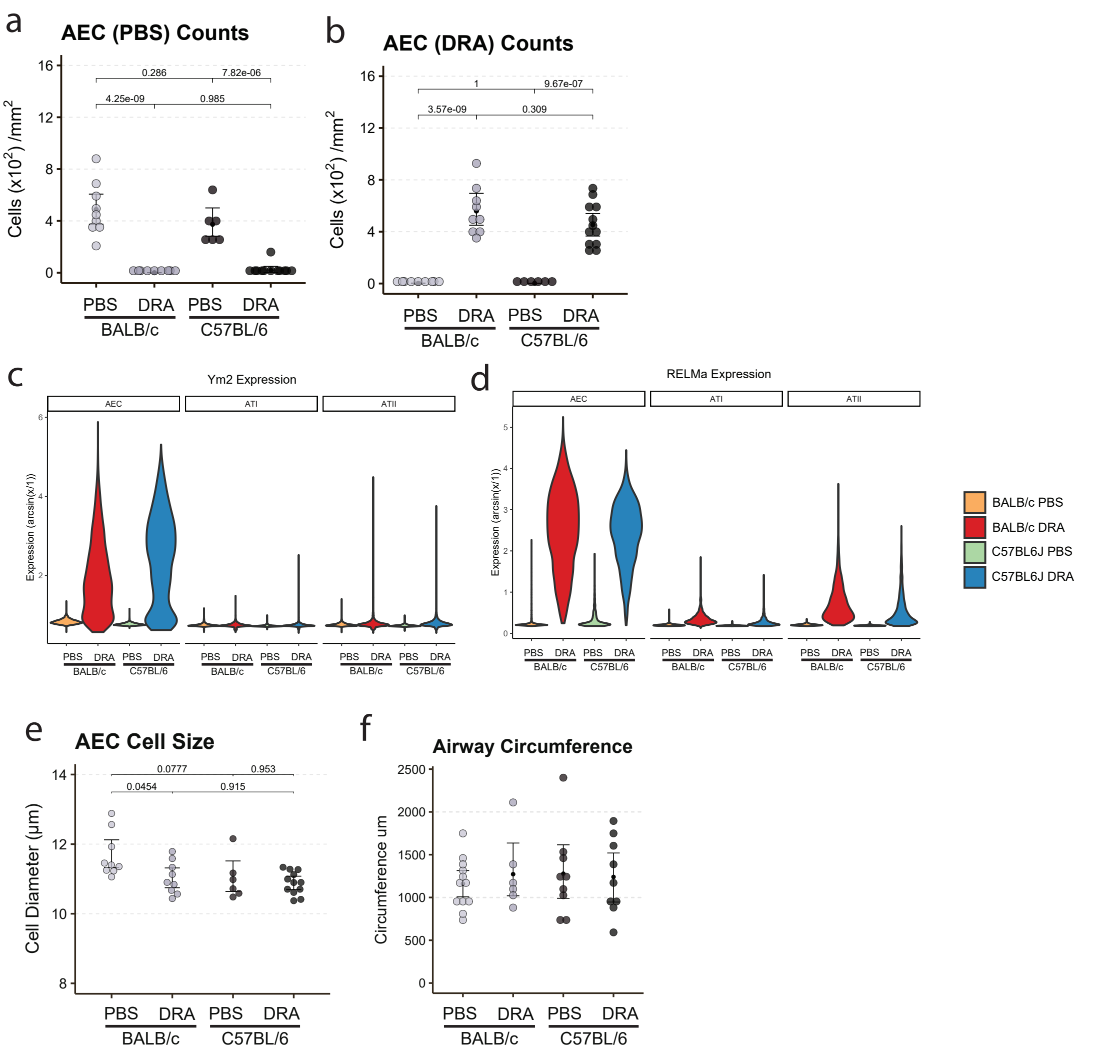


Supplementary Figure 6

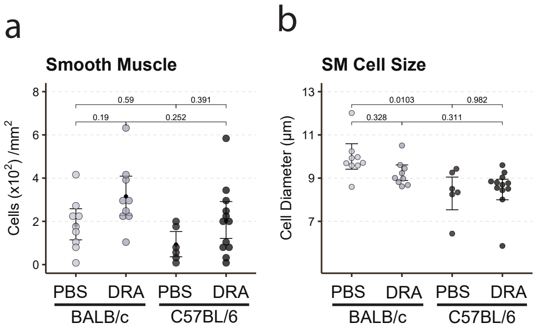


Supplementary Figure 7
